# Aneuploidy promotes transient stress adaptation and metabolic flexibility in the human fungal pathogen *Aspergillus fumigatus*

**DOI:** 10.1101/2025.09.29.679244

**Authors:** Anna E. Lehmann, Enrique Aguilar Ramírez, Nancy P. Keller, Joseph Heitman

## Abstract

Aneuploidy causes genome plasticity and enables adaptive responses that increase stress resistance and facilitate persistence under changing conditions in eukaryotes ranging from fungal pathogens to human cancer cells. *Aspergillus fumigatus* is a soil-resident mold and the most prevalent etiologic agent of invasive fungal infections globally. Invasive aspergillosis has an alarmingly high mortality rate, and fungal persistence in the infection microenvironment often renders treatments unsuccessful. Treatment failures can result from innate or acquired antifungal drug resistance and from fungal adaptation to other stressors encountered during infection including nutrient limitation, immune cell activity, and changes in pH and oxygen tension. Survival under changing environmental and host conditions therefore necessitates an ability to dynamically adapt to stress. We find that exposure of *A. fumigatus* to FK506, an antifungal and immunosuppressive *Streptomyces* natural product that inhibits the master regulatory phosphatase calcineurin, selects for unstable whole-chromosome aneuploidies that alleviate the polarized growth defects caused by calcineurin inhibition. Transcriptomic analysis revealed that Chr7 disomy leads to induction of the normally-silent neosartoricin biosynthetic gene cluster in response to FK506 exposure, and we demonstrate that constitutive genetic induction of this cluster is sufficient to largely recapitulate in a haploid background the response to FK506 in the aneuploid state. We further show that aneuploids undergo extensive metabolic rewiring but do not produce detectable neosartoricin, revealing global and cross-pathway effects resulting from both the aneuploid state and activation of *nscR*. We also demonstrate that the aneuploid chromosome duplications reduce susceptibility to the clinical antifungal voriconazole, underscoring the utility of aneuploidy as a flexible adaptive strategy. These findings represent a major advance in understanding how aneuploidy transiently alters stress responses and metabolism in the deadly human pathogen *A. fumigatus*.

**Highlights:** - *A. fumigatus* rapidly gains and loses supernumerary chromosomes in response to the calcineurin inhibitor FK506.

- Aneuploidy leads to global transcriptional and metabolic rewiring.

- Overexpression of the secondary metabolite gene cluster regulator *nscR* is sufficient to largely recapitulate the aneuploid response to FK506 in a haploid background.

- Chr4, 6, and 7 aneuploidies reduce susceptibility to voriconazole without altering expression of the azole target gene *cyp51A*.

## Introduction

Aneuploidy, an abnormal complement of whole chromosomes or chromosome segments, is a conserved adaptive strategy across eukaryotes. Whole-chromosome and segmental aneuploidies are ubiquitous in human cancers and are hypothesized to enhance tumor survival and proliferation in a dynamic and context-specific manner by altering gene dosage and activating generalized stress responses (reviewed in^1–3^). Aneuploidy is also an important persistence strategy in single-celled eukaryotes. In the model yeast *Saccharomyces cerevisiae*, unstable aneuploidy can serve as a rapidly-acquired, temporary solution to diverse stressors that potentially provides an avenue to less deleterious, stable genetic change^4–8^. Aneuploidy is also known to underlie unstable stress adaptation in multiple human and plant fungal pathogens^9^. Significantly, aneuploidy can serve as a major cause of reduced susceptibility to triazoles, the primary antifungal class administered clinically. In *Candida albicans*, azole resistance results from isochromosome formation of the left arm of Chr5, which harbors the genes that encode the azole target enzyme Erg11 and the drug efflux pump-activating transcription factor Tac1^10–12^. Whole-chromosome aneuploidy has also been reported to reduce *in vitro* antifungal susceptibility in the pathogenic yeast *Cryptococcus neoformans*^13,14^ and the plant pathogen *Aspergillus flavus*^15^. Additionally, a recent report identified segmental duplications of regions encoding the azole target-encoding genes *cyp51A* and *cyp51B* in azole-resistant *Aspergillus fumigatus* clinical isolates, revealing a correlation between copy number variation and clinically-relevant antifungal resistance in this globally ubiquitous pathogen^16^. These observations support a broadly conserved role for aneuploidy in stress adaptation in pathogenic fungi. Despite the apparent conservation of this phenomenon, however, genomic plasticity as an unstable adaptive mechanism remains underexplored in filamentous fungi.

*A. fumigatus* is a globally-distributed, soil-resident fungus and the leading cause of invasive fungal infections worldwide^17^. This organism has evolved a range of strategies that facilitate persistence in its harsh and variable ecological niches, and this same suite of adaptive traits can contribute to its pathogenicity. One such adaptation is the inducible production of secondary metabolites, two of the most well-studied being 1) DHN melanin, which protects conidia from UV radiation and reactive oxygen species and inhibits host cell apoptosis^18–20^ and 2) gliotoxin, which is toxic to mammalian host cells^21–24^. Numerous other secondary metabolites exhibit both antimicrobial activity against soil microbes and cytotoxic activity against host immune cells, suggesting context-dependent and evolutionarily-relevant reasons for their maintenance in the genome and selective induction (reviewed in^25,26^). The genes involved in secondary metabolite biosynthesis are encoded in self-contained and spatially discrete biosynthetic gene clusters (BGCs), which are often transcriptionally inactive under standard laboratory culture conditions^27^. Difficulties in activating silent BGCs continue to hinder efforts to elucidate the functions of these enzymes and products in fungal physiology and adaptation. Recent efforts to activate normally-silent BGCs have employed co-culture with soil microbes including *Streptomyces* and other Aspergilli that co-occupy a saprophytic environmental niche^28–30^. Several such studies have resulted in the identification of new natural products, suggesting a role for these compounds in cross-talk and interaction with other, potentially competing microbes and further evidencing their ecological relevance^28,29,31–34^.

Beyond the triggers for transcriptional activation being largely unknown, BGC induction is further complicated by the often-observed endogenous crosstalk of metabolites within a species. Myriad examples of this cross-talk have been observed in *A. fumigatus*, including loss of one metabolite (e.g. fumigaclavine C or hexadehydroastechrome) resulting in increased production of another metabolite (fumitremorgins)^35,36^, evidence of an enzyme from one BGC active in synthetic potential in another BGC^37^, and changes in BGC gene expression upon deletion of genes in a different BGC^38^. This complexity underscores the intricate and fine-tuned biology underlying *A. fumigatus* secondary metabolism and the wealth of information that remains to be uncovered about these processes.

Despite the apparent ubiquity of aneuploidy as a rapidly-acquired, flexible adaptive mechanism in eukaryotes, the potential roles of duplication and loss of genetic material in *A. fumigatus* stress responses remains a significant gap in knowledge. Here, we identify whole-chromosome aneuploidies in the *A. fumigatus* reference strain A1163 and in several clinical and environmental isolates. We demonstrate that Chr7 disomy induces transcription of the neosartoricin (*nsc*) BGC, which is silent under standard laboratory culture conditions. Induction of the *nsc* gene cluster transcription factor *nscR* in the A1163 haploid background (*nscR^OE^*) is sufficient to alter phenotypic responses to FK506 in a manner consistent with that observed in Chr7 aneuploids. Neosartoricin is produced by *nscR^OE^*, but not Chr7 aneuploids, suggesting a potential additional role for NscR in stress responses and/or metabolism. *nscR^OE^* and Chr7 aneuploids also undergo broad changes in both primary and secondary metabolism, and these expression changes do not necessarily correspond to expression changes in synthetic pathway genes. These findings reveal that, in addition to altering the transcriptome, aneuploidy also functions to promote metabolic flexibility. We also find that FK506-selected aneuploid chromosome duplications reduce susceptibility to the clinical antifungal voriconazole, illustrating the capacity for aneuploidy to confer fitness advantages under distinct and disease-relevant conditions. These findings extend evidence of whole-chromosome aneuploidy as a conserved adaptive strategy to the important fungal pathogen *A. fumigatus*, revealing a novel avenue by which dynamic stress adaptation and metabolic rewiring occur in this species.

## Results

### Unstable aneuploidy alleviates hyphal growth defects caused by FK506 in the A. fumigatus reference strain A1163

Our previous work has utilized FK506, an antimicrobial and immunosuppressive compound produced by *Streptomyces*, to uncover stable and unstable genetic adaptive strategies in the fungal pathogens *Mucor*^39,40^ and *Cryptococcus*^41–43^. FK506 binds to the endogenously-expressed protein FKBP12 and forms a complex that inhibits the protein phosphatase calcineurin, a mechanism that is broadly conserved in eukaryotes^44–46^. In *A. fumigatus*, calcineurin inhibition by pharmacologic or genetic means impacts a host of processes including development and morphology of conidia, septation, polarized growth, and cell wall composition^47–51^. The result of these severe growth and morphological defects is an abolition of virulence in pathogenesis models, underscoring the essentiality of calcineurin-mediated processes to the pathogenic capacity of this organism^48^.

While several downstream effectors of calcineurin have been described, most extensively the transcription factor CrzA^49,52,53^, detailed mechanistic bases for calcineurin-dependent morphological traits remain ill-defined. To further delineate these important processes, we employed a suppressor screen with the goal of elucidating genetic and/or epigenetic means by which *A. fumigatus* adapts to calcineurin inhibition. Conidia were plated on *Aspergillus* minimal medium (AMM) containing FK506 at a concentration of 1 µg/mL, which severely impairs growth of the A1163 wild-type strain (Fig. 1*A*). Colonies that exhibited increased vegetative growth were isolated and single-spored. A total of 52 FK506-adapted isolates were obtained, and whole-genome sequencing revealed medium- or high-impact mutations in 14 isolates, occurring in 6 genes that we predict are involved in bypass suppression of calcineurin (Fig 1*B*; Table S1). Surprisingly, a majority (34/52) of these isolates had duplications of 1 or more full chromosomes, and none of these aneuploids had high- or medium-impact sequence changes that we predict are likely to impact the observed phenotypic changes (Fig 1*B*; Table S1). We interestingly did not identify any mutations in the genes encoding the FK506 target, *fkbp12-1*, or the calcineurin catalytic or regulatory subunits, *cnaA* and *cnaB*. One possible reason for the failure to isolate mutants in these genes could be due to aneuploidy conferring a lower fitness cost than these genetic mutations, and/or the relatively short timeframe and growth-permissive concentration of drug used.

**Figure 1.**
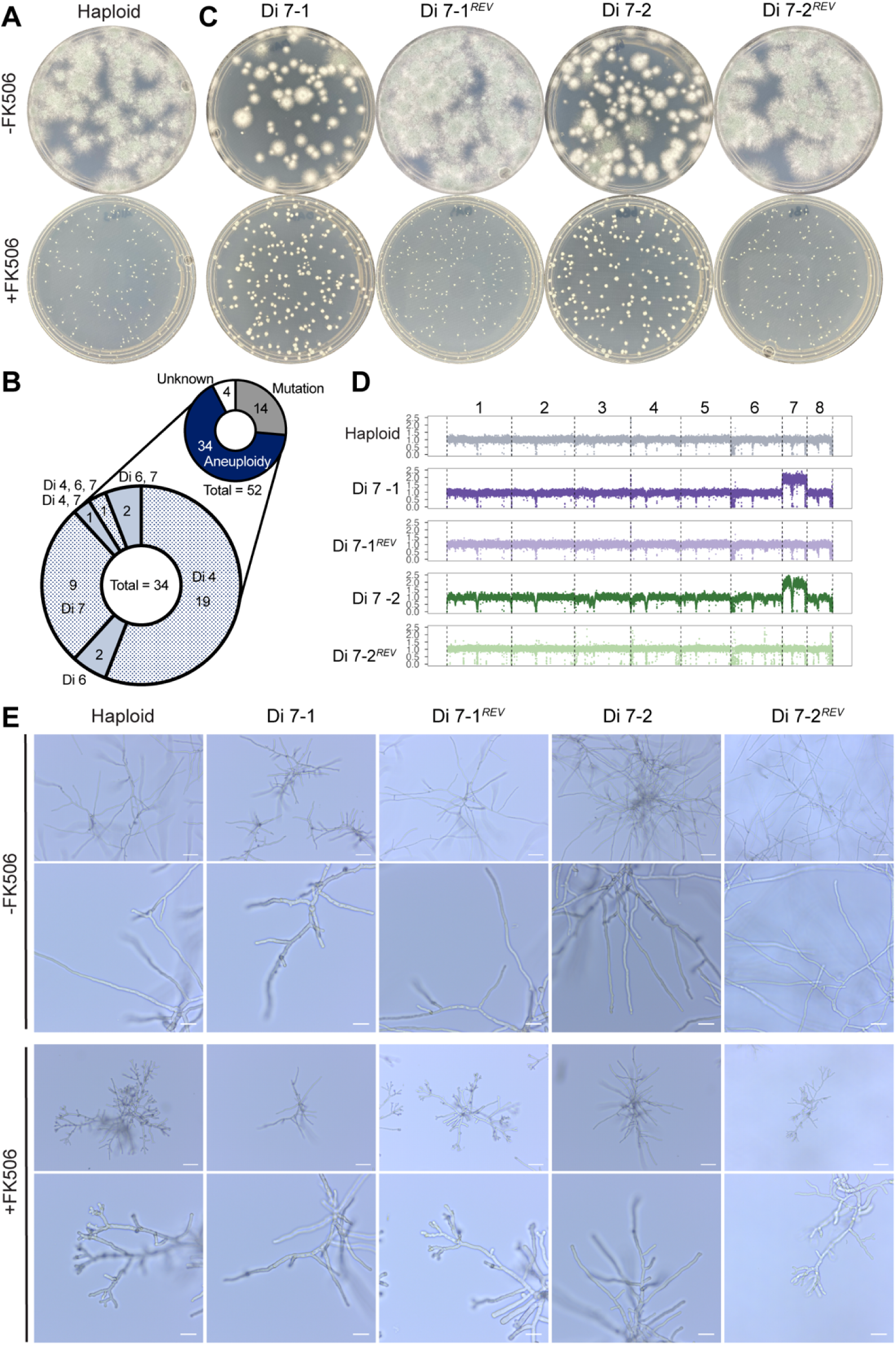
Unstable aneuploidy alleviates polarized growth abnormalities caused by FK506. (*A*) A1163 WT growth on AMM – (top) or + (bottom) 1 µg/mL FK506 for 3 days at 37°C. (*B*) Distribution of aneuploidies, high- or medium- impact mutations, and unknown mode of change determined from analysis of WGS data from FK506-adapted isolates. Enlarged pie chart shows frequency of chromosome duplications in the set of 34 aneuploids. (*C*) 2 representative Chr7 aneuploids grown on AMM + 1 µg/mL FK506 for 3 days before (top) and after (bottom) passage on drug-free AMM. Representative images from n ≥ 3 biologically independent replicates. (*D*) Normalized read depth averaged over 500 bp intervals, plotted relative to the closely-related CEA10 reference assembly, for strains in (*C*). (*E*) Conidia were inoculated in AMM +/- 1 µg/mL FK506 in a microscopy growth chamber and grown at 37°C for 24 hours. In each panel (-FK506 and +FK506), scale bars indicates 50 µm (top) and 20 µm (bottom). Representative images from n ≥ 3 biological replicates.

We elected to first focus our study on Chr7 aneuploids due to their prevalence in our screen (9/52 isolates) and the fact that Chr7 is the smallest chromosome in the genome, and previous work in yeast, *Drosophila*, and human cells has demonstrated that aneuploidies of smaller genomic regions are often better tolerated than those of larger regions^3^. Chr7 disomic strains (Di 7) have a growth defect on medium without FK506 (Fig. 1*C*), and when Di 7 conidia were plated on drug-free medium we observed heterogeneity in colony morphology that we hypothesized could be due to heterogeneous gain or loss of supernumerary chromosomes. To assess aneuploid chromosome stability in the absence of FK506, strains were cultured on drug-free AMM for at least 9 days. After drug-free passage, conidia were re-plated on FK506 medium. Passage on drug-free medium resulted in reversion to a wild-type, haploid level of growth on FK506 (Fig. 1*C*). Sequencing of the reverted (*REV*) isolates showed a corresponding loss of the aneuploid chromosomes, linking the observed morphological changes directly to the gain of a second copy of Chr7 (Fig 1*D*; Table S1).

At the beginning of this study, we identified 6 isolates that originated from a single initial pool of conidia but formed separate colonies with similar enhanced growth phenotypes on individual FK506 selection plates. Whole-genome sequencing of these isolates revealed that all 6 carried an extra copy of Chr7 and differed from the progenitor strain by several identical SNPs in coding regions of the genome (Table S1). Based on the genetic and ploidy similarities, we hypothesize that these isolates may have originated from a clonal population of Di 7 conidia in the initial pool. Due to their apparent clonality, these 6 isolates are treated as a single independent event (Di 7-1) in all results presented here. To avoid identifying additional clonal isolates, and to confirm that the additional SNPs present in the aneuploids are not responsible for the aneuploid karyotype, all other isolates described in this study were isolated from 20 different single-spore derived populations of conidia from the original A1163 stock, and one FK506-adapted colony was isolated per replicate.

Consistent with previously-described effects of calcineurin inhibition^48,50,54^, and stress responses more generally^55,56^, we observed defects in polarized growth, indicated by increased density of hyphae, stunted extension, and aberrant branching at the apical hyphal tips, in the haploid +FK506 (Fig. 1*F*). In contrast, Chr7 aneuploids lack the hyper-branched hyphal tips observed in the haploid, and these phenotypes are reversed to a state that is indistinguishable from the haploid progenitor in the reverted (*REV*), haploid strains derived from the Chr7 aneuploids (Fig. 1*F*). These results further implicate Chr7 aneuploidy in the reversal of polarized growth abnormalities caused by calcineurin inhibition and indicate that aneuploidy leads to an improved ability to tolerate stress caused by FK506.

### Aneuploidy enhances growth on FK506 in distinct A. fumigatus isolates

Unstable adaptive responses to antifungal compounds are ill-defined in *A. fumigatus*, and we questioned whether the high prevalence of unstable aneuploidy in response to FK506 occurred due to a specific property of A1163, or if this transient response could be generalized to other isolates. We selected 10 diverse, non-reference strains representing the 3 clades of *A. fumigatus*^57^, as well as both clinical and environmental sources (Table S2). 3 of these strains exhibited robust growth on FK506 at concentrations up to 50 µg/mL (Fig. S1*A*). Of the remaining 7 strains, we successfully repeated the screen used for A1163 at 1 µg/mL FK506 for 6 strains and 0.5 µg/mL for 1 strain. 3 to 4 independent isolates with improved growth on FK506 were obtained per strain for a total of 26 isolates. All isolates were passaged on drug-free medium for at least 15 days, and re-plating conidia on FK506 after drug-free passage showed a loss of the growth improvements initially observed in 12 of these isolates (Fig. S1*B*). In our relatively small sample size, the number of stable and unstable responses was not evenly distributed between strains, with some giving rise to only stable isolates, some only unstable, and some a mix of the two (Fig. 2*A*). Nevertheless, in total we obtained stable and unstable isolates in roughly equal proportions (14 stable, 12 unstable). Sequencing of the unstable FK506-adapted isolates revealed, similar to A1163, widespread aneuploidy involving disomies of Chr4 and Chr7 (Fig. 2*B*). While all aneuploidies were unstable, and no stable isolates were aneuploid, not all unstable isolates had detectable aneuploidies (Fig. 2*B*; Fig. S2). Because aneuploids are highly unstable, one possibility is that the non-aneuploid unstable isolates were aneuploid upon isolation and lost the duplicated genetic material during culture prior to DNA extraction. Another possibility is that a non-aneuploidy unstable adaptive mechanism, such as chromatin remodeling or RNA interference-mediated gene silencing, is responsible for the FK506 responses observed in these isolates^39,58^. Together, these results confirm that unstable aneuploidy is not restricted to the A1163 reference strain, can occur frequently under the proper selective conditions, and is conserved in strains from clinical and environmental sources representing at least clades 1 and 2.

**Figure 2.**
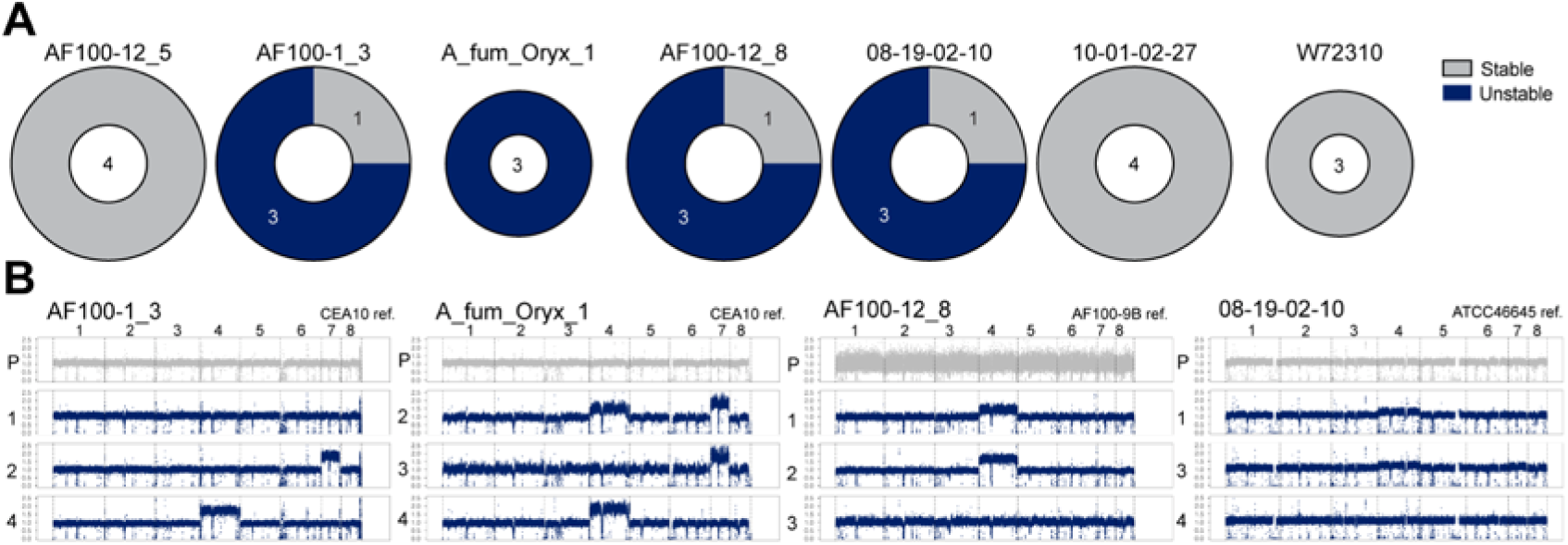
Stable and unstable FK506-adapted isolates arise in diverse strain backgrounds. Conidia from clinical and environmental strains were plated on AMM +0.5 µg/mL FK506 (W72310) or 1 µg/mL FK506 (all other strains) and colonies with improved vegetative growth were selected. Stability was determined by passaging isolates on drug-free AMM for at least 15 days and re-plating on FK506. (*A*) Distribution of stable and unstable FK506-adapted isolated identified in each strain background. (*B*) Normalized read depth for progenitor strain (P) and unstable FK506-adapted isolates that arose in 4 strain backgrounds, plotted relative to a closely-related assembly as indicated at the top right of each plot.

### Chr7 disomy induces expression of secondary metabolite biosynthetic genes

Aneuploidy in yeasts and humans is known to alter gene expression and phenotypic traits both through dosage-dependent transcriptional changes and by activating dosage-independent, aneuploidy-specific responses. In *Candida albicans*, i5L isochromosome formation confers fluconazole resistance by altering copy number and expression of *ERG11*, which encodes the azole drug target enzyme, and *TAC1*, which encodes an efflux pump-activating transcription factor^59^. We therefore hypothesized that changes in gene expression in Chr7 aneuploids resulting from the changes in copy number and/or the response to altered DNA content could explain the phenotypic changes observed in the A1163 Chr7 aneuploids. To probe a potential transcriptional basis for the observed phenotypic changes, we conducted 2 independent RNA sequencing experiments with the haploid progenitor strain and independent Chr7 aneuploids (Di 7-1 and Di 7-2), with and without FK506. Total RNA was extracted from triplicate cultures and analyzed by paired-end bulk mRNA sequencing (Materials and Methods). Parallel DNA sequencing from Di 7-1 cultured under the drug-free RNA extraction condition confirmed that the chromosome duplication was maintained in the drug-free condition (Fig. S3*A*).

RNA-seq reads were mapped to the A1163 reference genome^60^, and PCA was performed analyzing the top 1000 variable transcripts in each experiment, which capture the majority of variance in the data (Fig. S3*B, C*). In both cases, the majority of the variance was explained in principal components 1 and 2 (Fig. S3*D, E*). PCA indicates that for Di 7-1, drug condition is the primary factor explaining gene expression differences (PC1; 52.2% of variance), followed by ploidy (PC2; 21.6% of variance) (Fig. S3*F*). In contrast, in Di 7-2, variance is primarily driven by ploidy (PC1; 60.5%), with drug condition explaining 19.3% of the observed variance (Fig. S3*G*). In both experiments, analysis of differentially expressed genes (DEGs; p_adj_ ≤ 0.05 and |log_2_FC| ≥ 1) between aneuploids and haploid +FK506 revealed large-scale changes in gene expression, with 291 upregulated and 167 downregulated genes in experiment 1 and 1537 upregulated and 442 downregulated genes in experiment 2 (Fig. 3*A, B*; Table S3). Similar to the PCA results, these differences in gene expression between the two experiments illustrate a much larger effect of aneuploidy on the global transcriptome in Di 7-2 than in Di 7-1.

**Figure 3.**
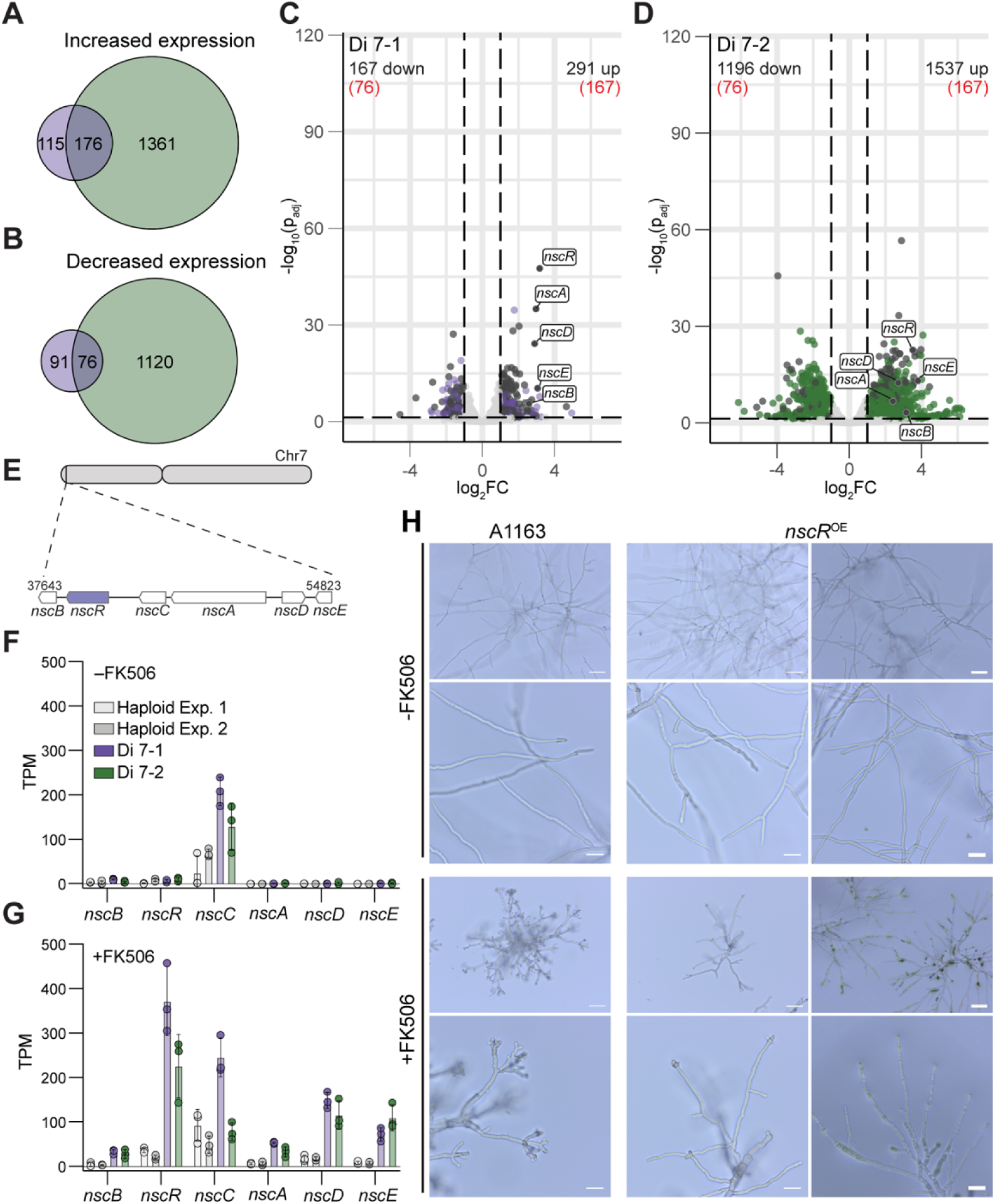
Secondary metabolite transcription factor *nscR* is induced in Chr7 aneuploids and recapitulates the Chr7 aneuploid FK506 response in the haploid. (*A* and *B*) Genes with increased (*A*) and decreased (*B*) expression in comparison between Chr7 aneuploid and haploid in 2 RNA-seq experiments. (*C* and *D*) Differentially expressed genes in comparison between Chr7 aneuploid +FK506 and WT +FK506 in experiment 1 (*C*) and 2 (*D*). Grey colored points denote shared up- and down-regulated genes between the two experiments. Dashed lines indicate log_2_FC = ±1 and p_adj_ = 0.05. (*E*) Location of the *nsc* BGC on Chr7. (*F* and *G*) Transcript abundance (TPM) of *nsc* BGC genes in haploid and Chr7 aneuploids -FK506 (*F*) and +FK506 (*G*). Points represent technical replicates; error bars indicate standard deviation. (H) Hyphal growth phenotypes of A1163 wild-type and *nscR^OE^*, +/- FK506, as in Fig. 1E.

We predicted that phenotypic commonalities in response to FK506 observed in Di 7-1 and Di 7-2 would most likely be explainable by transcriptional change(s) shared between the two strains. Comparison of the up- and down-regulated gene sets between the two experiments showed that Di 7-1 had 176 up-regulated and 76 down-regulated genes in common in the FK506 condition (Fig. 3*A*, *B*). Interestingly, 4 genes in the *nsc* biosynthetic gene cluster (*nsc* BGC), which regulates synthesis of the secondary metabolite neosartoricin^61^, were among the most highly induced genes in both experiments (Fig. 3*C*, *D*; Table S3). The *nsc* BGC, located near the left telomere of Chr7, encodes the transcriptional regulator *nscR* and the neosartoricin biosynthetic enzymes *nscA-E* (Fig. 3*E*). This gene cluster, with the exception of the FAD-dependent monooxygenase *nscC*, is silent or lowly expressed in the haploid and Chr7 aneuploids -FK506, and minimally induced in the euploid +FK506 (Fig. 3*F, G*). FK506 exposure induces *nsc* gene expression specifically in the aneuploids, with *nscR* having the highest expression (Fig. 3*G*).

Secondary metabolite BGCs can be transcriptionally activated by both pathway specific transcription factors and global regulators or epigenetic modifications that themselves are induced by various environmental stimuli^62–64^. Given the large number of transcriptional changes observed in the Chr7 aneuploids, we assessed whether aneuploidy globally alters secondary metabolite BGC transcription or whether the effect is specific to the *nsc* BGC. TPM quantification of the genes in the 18 additional known A1163 secondary metabolite BGCs^65^ shows that these clusters are either silent in all conditions, or do not undergo the same cluster-wide differential change as the *nsc* BGC in response to aneuploidy and FK506 (Fig. S4). One such example is the fumagillin BGC monooxygenase *fmaE* (AFUB_086100), which is constitutively highly expressed across strains and conditions while the other genes in its cluster are not expressed. Several genes in the hexadehydroastechrome and trypacidin BGCs also undergo increased transcription in response to FK506. Unlike the *nsc* BGC, however, most genes in these clusters remained silent. These results led us to conclude that Chr7 duplication leads to specific induction of the *nsc* BGC expression, rather than general induction of secondary metabolic genes, in response to FK506 exposure.

### nscR overexpression recapitulates Chr7 aneuploid phenotypes in response to FK506

To directly explore a potential role for the *nsc* BGC in the aneuploid phenotypes observed in response to FK506, we generated a strain expressing *nscR* (AFUB_086680) under control of the *A. fumigatus* Af293 *gpdA* promoter in the A1163 euploid background (*nscR^OE^*). The *nscR* transcript is constitutively overexpressed 107-fold without FK506 and 91-fold with FK506 in this strain (Fig. S5*A*). In static submerged culture with FK506, *nscR^OE^* exhibits decreased hyphal hyperbranching similar to Chr7 aneuploids (Fig. 3*H*). Additionally, *nscR^OE^* exhibits a comparable radial growth defect to Chr7 aneuploids on drug-free medium, and a comparable, minor but statistically significant, increase in radial growth on medium containing FK506 (Fig. S5*B-D*). Taken together, these observations indicate that *nscR* overexpression specifically alleviates FK506-dependent polarized growth defects in a manner similar to Chr7 aneuploidy. Our findings thus reveal that overexpression of a single gene is sufficient to largely recapitulate a particular aneuploid stress response in a haploid background.

### Aneuploid secondary metabolite profiles distinguish them from haploid and nscR^OE^

We hypothesized that NscR may influence the response to FK506 through the production of neosartoricin, which could act as a signaling molecule. Alternatively, NscR might activate enzymes involved in other synthetic processes, or an enzyme that can degrade FK506. NscR could also have transcriptional activity outside of the *nsc* BGC that might influence the FK506 response. To investigate a potential correlation between *nscR* expression and neosartoricin production in Chr7 aneuploids and *nscR^OE^*, we profiled the metabolites produced in these strains by LC-MS. Surprisingly, we found that only *nscR^OE^* produces neosartoricin, while this compound is absent or at undetectable levels in the Chr7 aneuploids (Fig. 4*A-C*). Considering the many instances of metabolite cross-talk in *A. fumigatus*, we next expanded our metabolomics analysis to assess the abundances of well-characterized *A. fumigatus* secondary metabolites in the A1163 haploid, *nscR^OE^*, and Di 7 strains. This analysis revealed clear increases in pyripyropene A, fumitremorgin C, pseurotin A, and fumagillin biosynthesis (Fig. 4*D*; Fig. S6). Of these metabolites, only fumitremorgin C showed a significantly increased abundance in both the Chr7 aneuploids and *nscR^OE^*(Fig. S6*A*). These changes in secondary metabolite production occurred in the absence of clear transcriptional induction of their BGCs, indicating potential non-concordance between BGC activation and associated secondary metabolite production in the Chr7 aneuploids (Fig. S7). Additionally, the extensive changes in metabolite abundances that occur as a consequence of overexpressing a single transcriptional regulator, *nscR*, underscore the complexity in regulation of these processes and the potential for perturbation of one biosynthetic pathway to exert widespread effects on secondary metabolism more broadly. We also examined the abundance of FK506 in haploid, Chr7 aneuploids, and *nscR^OE^* and observed no significant difference in its abundance in the Chr7 aneuploids and a small, but statistically significant, reduction in abundance in *nscR^OE^* (Fig. S6*E*). Despite this reduction in FK506 abundance, the level of the compound remained relatively high in all of the samples, which led us to conclude that elimination of FK506 alone is unlikely to explain the shared FK506 responses.

**Figure 4.**
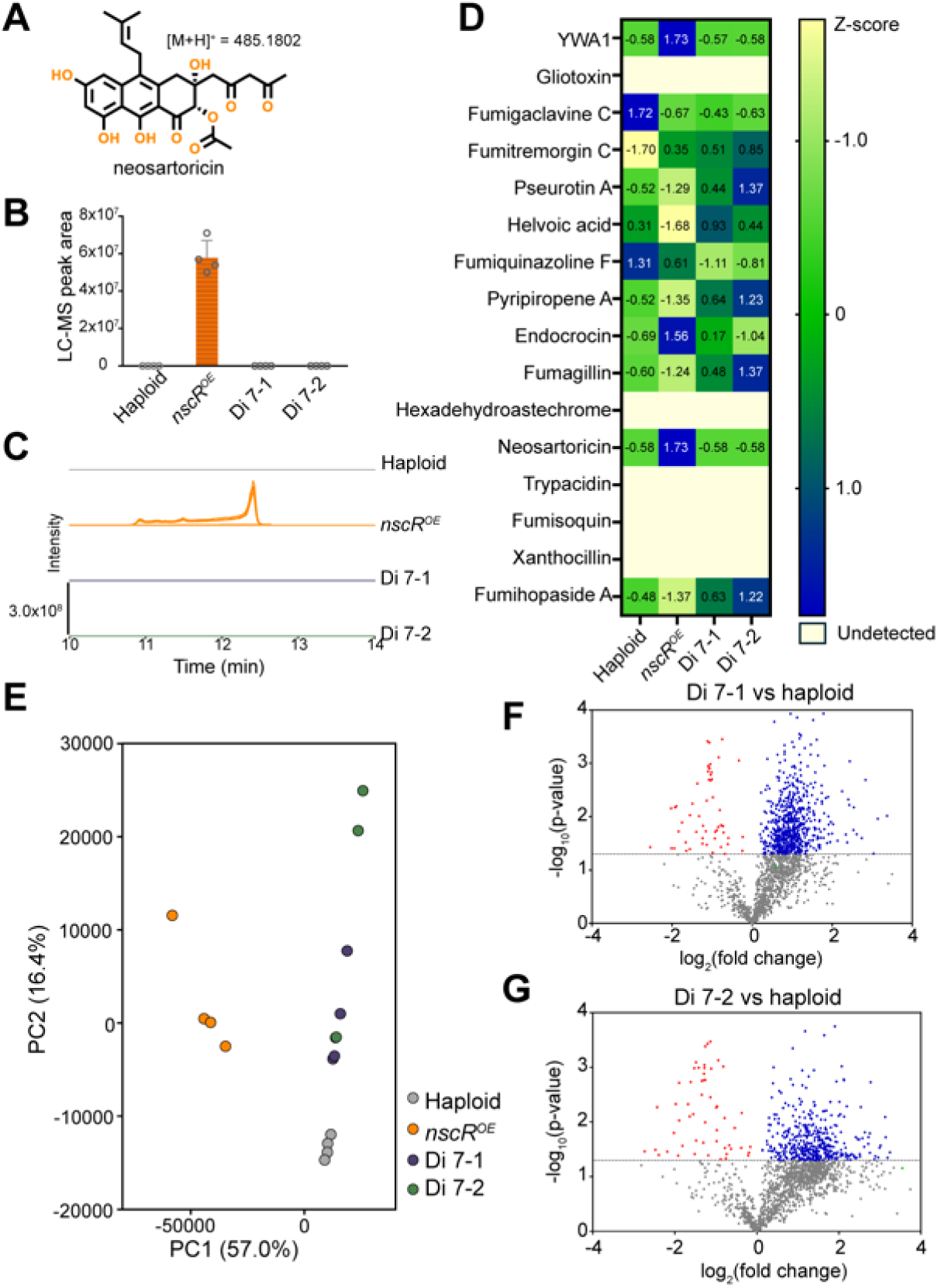
Chr7 aneuploids and *nscR^OE^* undergo extensive metabolic rewiring. (*A*) Chemical structure of neosartoricin. (*B*) Abundance of neosartoricin in the different strains, determined based on the average area under the curve of the detected peak for the molecular ion [M+H]+ = 485.1802 across the four replicates analyzed, and (*C*) its associated chromatogram. (*D*) Heatmap representing the abundance of well-characterized secondary metabolites of *A. fumigatus*, based on the average area under the curve associated with their LC-MS features. (*E*) Principal component analysis (PCA) showing the variance among the different strains according to the abundance of all LC-MS–detected features in positive mode. (*F* and *G*) Volcano plots comparing the abundance of detected features (in positive mode) between the aneuploids Di 7-1 (*F*) and Di 7-2 (*G*) and the haploid. The feature corresponding to fumitremorgin C is highlighted in green. Four replicates are included in all representations.

The relative abundances of all LC-MS features associated with *A. fumigatus* metabolites were next compared across strains by principal component analysis (PCA). The resulting chemical profiles segregated into three distinct groups: haploid, *nscR^OE^*, and the Di 7 strains, indicating that, at a global level, the Chr7 aneuploids and *nscR^OE^* display chemical profiles clearly distinct from one another and from the haploid wild-type (Fig. 4*E*). To further elucidate how the Chr7 aneuploids differ from their haploid progenitor, we conducted a volcano plot analysis using the LC-MS features, which revealed that a large number of metabolites were overproduced in the aneuploids relative to haploid (Fig. 4*F*, *G*). Interestingly, many of these metabolites were consistent with ergosterol-type triterpenes, based on their fragmentation patterns and predicted formulas (Fig. S8). These dramatic alterations in metabolism, coupled with the non-concordance between transcription and metabolites produced, indicate that aneuploidy of Chr7 promotes widespread metabolic rewiring, and that these changes are incompletely explained by changes in gene expression.

### Chr4, 6, and 7 aneuploidies reduce voriconazole susceptibility

Azole antifungals are direct inhibitors of the ergosterol biosynthetic enzyme Cyp51A, and Cyp51B is a parallel biosynthetic enzyme that is not a primary target of the drug but has been proposed to have a role in compensating for loss of Cyp51A activity in azole-resistant isolates^66^. Increased expression of *cyp51A* via promoter mutations is an established azole resistance mechanism in *A. fumigatus*^67–73^, and segmental duplications of genomic regions encoding *cyp51A* (Chr4), *cyp51B* (Chr7), and the *cyp51* electron carrier protein *cyp51ec* (Chr5) were recently correlated with elevated azole resistance in *A. fumigatus* clinical isolates^16^. *cyp51A* copy number increases in these isolates correspond to increases in *cyp51A* transcript abundance under itraconazole treatment, leading to the hypothesis that the increase in gene dosage of the drug target leads to an increased concentration of drug needed to inhibit fungal growth^16^. These observations are consistent with an established azole resistance mechanism in *C. albicans*, in which duplication of Chr5L leads to fluconazole resistance by increasing the copy number of the azole target-encoding gene *ERG11*^10,59^.

Given the presence of Chr4 and Chr7 disomies in our dataset, along with the observed changes in ergosterol biosynthesis, we investigated whether aneuploidy affects susceptibility to voriconazole. We measured the MICs of one aneuploid representing each karyotype observed and its corresponding reverted (*REV*) isolate (Fig. 5*A*). We found that aneuploids with duplications of Chr4, Chr6+7, and Chr4+6+7 have significantly higher MICs than haploid (Fig. 5*B*). The MICs of the *REV* isolates were not significantly different than those of the haploid progenitor. We hypothesized that the decreased voriconazole susceptibility in aneuploids may be due to an increase in *cyp51A* and/or *cyp51B* transcript abundance. We measured *cyp51A* and *cyp51B* transcript abundances by qPCR and observed no significant differences in *cyp51A* or *cyp51B* transcript abundances in any of the aneuploids relative to the haploid (Fig. 5*C-F*). To ensure that this was not due to some property of the specific culture conditions or medium used, we performed the experiment utilizing an additional culture condition (shaking flask cultures in AMM with DMSO or 1 µg/mL voriconazole) and again observed no changes in transcript abundance of the azole target-encoding genes (Fig. S9). These observations strongly suggest that, in contrast to the known aneuploidy-driven resistance mechanism in *C. albicans*, azole susceptibility in the *A. fumigatus* whole-chromosome aneuploids is not associated with an increase in copy number or expression of the drug target genes under drug treatment. Future work should investigate the potential metabolic, transcriptomic, and/or proteomic signatures in aneuploids under azole treatment to shed light on the means by which transient supernumerary chromosome acquisition can enhance tolerance of this important clinical antifungal.

**Figure 5.**
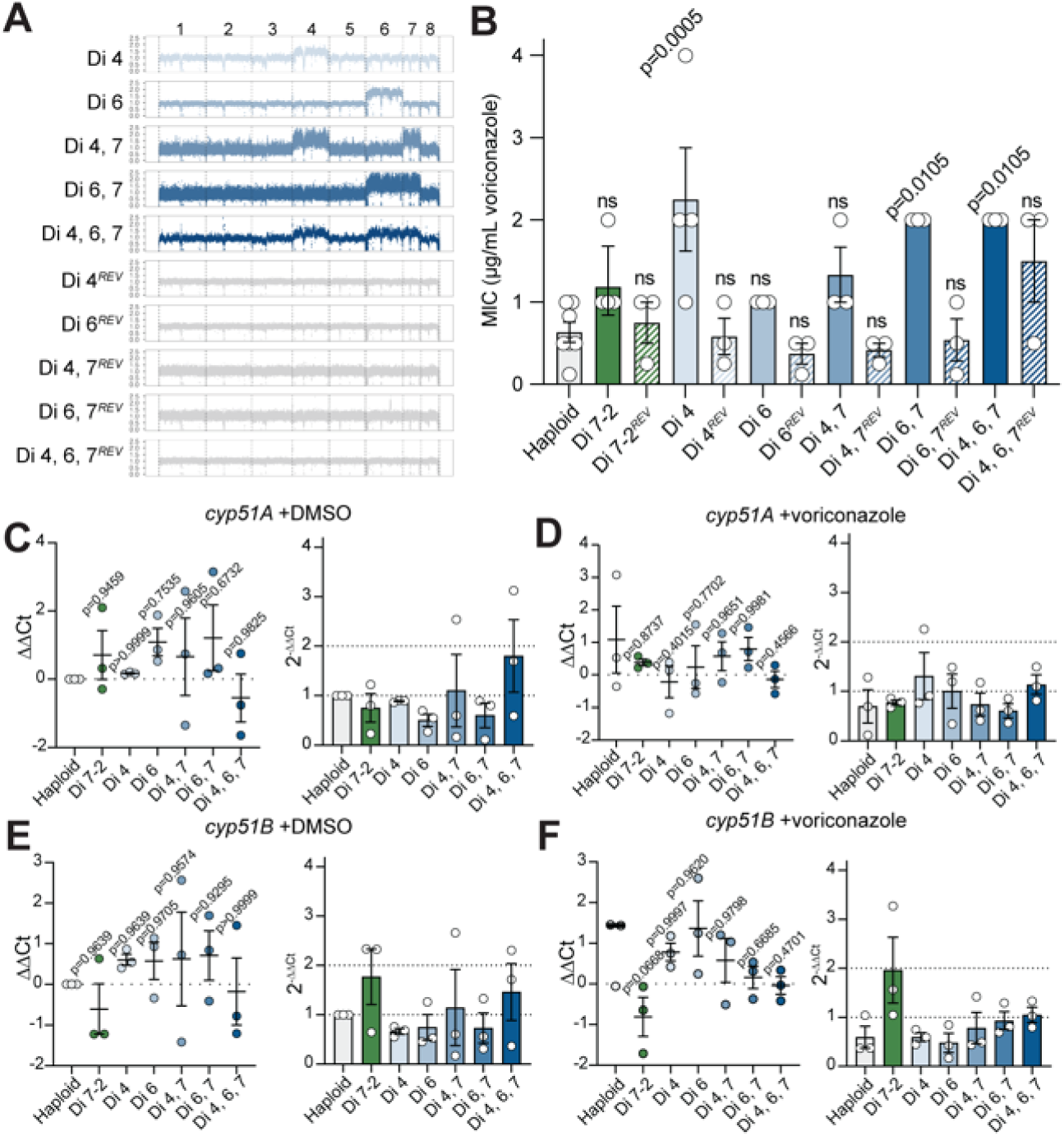
Chr4, 6, and 7 aneuploidies reduce voriconazole susceptibility without changes in *cyp51A* or *cyp51B* gene expression. (*A*) WGS coverage plots showing different karyotypes tested. (*B*) MICs (µg/mL voriconazole) measured by CLSI broth dilution; points represent biological replicates and statistical test is a one-way ANOVA with corrected Bartlett’s test. (*C-F*) Haploid and aneuploid strains were cultured in RPMI for 22 hours, then medium was removed and replaced with RPMI +DMSO or RPMI +0.2 µg/mL voriconazole. Fold change of *cyp51A* (*C, D*) and *cyp51B* (*E, F*) transcripts under DMSO (*C, E*) or voriconazole (*D, F*) treatment. Statistical significance was determined by one-way ANOVA with Dunnett’s multiple comparison test on ΔΔCt values. Fold change values, relative to haploid +DMSO, are represented as linearized ΔΔCt values (2^-ΔΔCt^).

## Discussion

Aneuploidy functions as a dynamic means of generating genomic and phenotypic diversity across the kingdom Eukarya, facilitating rapid stress adaptation in a genetically transient and readily-reversible manner. Our findings extend knowledge of this phenomenon to the important fungal pathogen *Aspergillus fumigatus*, revealing a function for whole-chromosome duplications in adapting to the calcineurin inhibitor FK506 and dynamically rewiring metabolism under FK506 stress. Aneuploidy is a hallmark of cancer, and metabolic reprogramming in aneuploid cancer cells is a major avenue supporting proliferation of tumor cells^74^. The Warburg effect, in which oncogenic cells shift to primary utilization of the oxygen-independent glycolytic pathway for energy metabolism irrespective of oxygen tension, is thought to promote tumor proliferation in part by increasing the rate of ATP production and supporting growth under hypoxia^75–78^. Glycolytic shifts in tumor cells have been linked to chromosome copy number changes, supporting a potential role for the ubiquitous aneuploidies found in cancer cells in influencing oncogenic metabolic transitions^79,80^. Similarly, in *S. cerevisiae* certain chromosomal aneuploidies increase mitochondrial respiration and lead to decreased pyruvate levels^81^. In *A. fumigatus*, metabolic plasticity is necessary for survival in nutrient-limited environments encountered during infection^82,83^. Metabolic flexibility is also critical for persistence in low-oxygen microenvironments generated during pulmonary infection and in drug-resistant biofilms, which become increasingly hypoxic as they mature^84,85^. Understanding the potential parallels in aneuploidy-driven metabolic rewiring in cancer cells, yeast, and *A. fumigatus* therefore has the potential to improve our understanding of this organism’s persistence in disease-relevant contexts.

Along with the A1163 reference strain, we isolated Chr4 and Chr7 aneuploids in 4 additional strain backgrounds from both clinical and environmental sources. Interestingly, 2 of the strains that readily gave rise to whole-chromosome aneuploids were isolated during a 4.5 year time course study from the sputum of a person with cystic fibrosis^86^. This finding provides direct evidence that there are isolates present in individuals with *Aspergillus* infections that are capable, under the correct selective conditions, of acquiring and maintaining unstable chromosome duplications. Another recent study, which identified multiple independent segmental duplications in azole-resistant clinical isolates^16^, further supports the hypothesis that genome instability is a generalizable and clinically-relevant phenomenon in *A. fumigatus*. Because aneuploidy appears to arise frequently under stress conditions across isolates, coupled with the observation that supernumerary chromosomes are rapidly lost in the absence of drug selection, it is highly plausible that prior sequencing efforts have failed to identify naturally-occurring aneuploids in clinical and environmental samples by culturing isolates under non-selective conditions prior to DNA-sequencing. With the knowledge that aneuploids are highly unstable, future efforts to detect transient karyotype diversity in natural populations can employ isolation strategies that maintain selective pressure. Such efforts will be important to understand the potential roles of aneuploidy in promoting clinically and agriculturally-relevant antifungal resistance and persistence under stress.

The majority of research on aneuploidy in pathogenic fungi to this point has focused on *Candida albicans*, where azole resistance arises as a consequence of increased copy number of *ERG11* and *TAC1*^10,59^. Recent findings by Khateb et al^16^ similarly showed that copy number duplication of the genomic region encoding *cyp51A* in azole-resistant *A. fumigatus* clinical isolates corresponds to increased transcription of the drug target gene, suggesting a parallel aneuploidy-driven resistance mechanism to that observed in *C. albicans*. In contrast to these findings, we showed that Chr4, 6, and 7 whole-chromosome aneuploidies decrease *in vitro* susceptibility to voriconazole, and do so without apparent changes in *cyp51* gene expression. Our findings suggest that important differences may exist in the mechanisms by which aneuploidy can alter azole susceptibility between different fungal pathogens and between different aneuploid isolates of the same species.

In this study, we investigated transcriptional changes in 2 independent Chr7 aneuploids under FK506 treatment and observed both shared and unique changes in gene expression between the strains. Because these two aneuploids exhibited phenotypic similarities under the tested stress condition, we focused on those differentially-expressed genes that were shared between them. Among this shared set of genes was the transcription factor *nscR*, and we demonstrated that overexpression of *nscR* in the haploid progenitor background largely recapitulates the FK506 stress response shared between the Chr7 aneuploids. However, the significant number of genes differentially expressed in one but not the other aneuploid could well underly differences in their responses to other stresses, and the consequences of these transcriptional differences warrant further investigation. The broad differences in transcription between two strains of the same genetic background and karyotype, along with the observation that the transcriptional changes were not specific to only those genes encoded on the duplicated chromosome, could indicate that there is some degree of non-specificity in aneuploidy-driven gene expression changes. Such non-specificity could serve as a bet-hedging strategy in which aneuploidy generates transient transcriptional diversity in an otherwise-clonal population, potentially increasing the odds of survival upon exposure to stress. Indeed, stochastic gene expression among clonal populations is known to underlie differences in competence in *Bacillus subtilis*^87^ and heterogeneous expression of stress response proteins in *S. cerevisiae*^88^. In *S. cerevisiae*, aneuploidy also leads to elevated non-genetic population heterogeneity and heterogeneous gene expression changes in response to heat shock and nutrient stresses^89^. These prior observations indicate that transcriptional heterogeneity between genetically identical aneuploids could have relevant consequences for fitness in environmental and infection environments, and that studying multiple aneuploids in parallel can aid in determining the particular aspects of the aneuploid response that are relevant to the stress condition in question.

We found that the normally-silent *nsc* BGC is specifically activated in aneuploids exposed to FK506, and that overexpression of the *nsc* cluster transcription factor *nscR* results in reversal of the polarized growth defects caused by FK506. These effects are independent of detectable neosartoricin production, suggesting that NscR may have functions outside of the production of this compound, and/or that a neosartoricin intermediate, rather than the final compound, could be the relevant molecule in the FK506 stress response. Ongoing work seeks to characterize potential additional functions of NscR in this context and their effects on stress responses and metabolism. Interestingly, deletion of the gene encoding the calcineurin-activated transcription factor *crzA* is also reported to increase *nsc* gene expression, but not neosartoricin production, further supporting a role for this gene cluster in relation to the calcineurin pathway^90^. Several additional studies have also shown that BGC transcript abundance does not always translate to concomitant metabolite production^91,92^. The mechanisms underlying non-concordance between transcript and metabolite production are understudied but can involve post-transcriptional and/or post-translational modifications. One such example is phosphorylation regulation of AflR, the pathway specific transcription factor required for sterigmatocystin and aflatoxin production^93^. Several studies have also demonstrated that transcription factors encoded within one BGC can regulate the expression of secondary metabolite genes in other BGCs, often those located on other chromosomes. For example, in *A. fumigatus*, HasA both positively and negatively regulates multiple BGCs^94^, in *Penicillium expansum*, XanC specifically activates the citrinin BGC^95^, and in *Aspergillus nidulans*, ScpR drives expression of both the *inp* BGC and the asperfuranone BGC^91^. Similarly, in *Fusarium graminearum*, TRI6 not only activates the trichothecene BGC but also influences genes involved in the isoprenoid/mevalonate pathway^96^. Our findings that aneuploidy broadly rewires metabolism, and that changes in secondary metabolite abundance do not necessarily correlate to transcriptional activation of their BGCs, build on this existing body of literature and provide further evidence for the complexity of secondary metabolism regulation. Further exploration of cross-talk and cross-regulatory functions of BGCs in aneuploids is likely to yield valuable insight into the mechanisms underlying metabolic flexibility and stress responses in this organism.

In the context of intra-population genetic diversity, increased copy number of full chromosomes may increase the odds of positive selection compared to stable genetic mutations by flexibly altering large sets of genes and rewiring metabolism in a readily-reversible manner. Given the variability in stress conditions encountered by *A. fumigatus* both in its environment and during host interactions, it is conceivable that aneuploidy could facilitate rapid adaptation and persistence in both ecological and disease contexts through transcriptional changes that can be turned on and off without the need for permanent genetic change. Overall, our results lay a foundation for future inquiry into how flexible genomic changes may drive beneficial adaptive responses and increase the odds of survival in response to stresses such as competing microorganisms, adverse conditions in the environment and during host infection, and exposure to clinical and agricultural antimicrobial compounds.

## Resource Availability

### Lead contact

Requests for resources and further information should be directed to, and will be fulfilled by, the lead contact, Joseph Heitman.

### Materials availability

All unique materials generated in this study are available without restriction from the lead contact.

### Data and code availability

All raw DNA- and RNA-seq data are deposited at NCBI under Bioproject PRJNA1335234, PRJNA1335235, and PRJNA1448995. Processed sequencing data (variant calls and DESeq results) are included in the Supplemental Material. All original code generated for this project is available at: https://github.com/annalehmann/Afum_FK506_resistance and https://github.com/annalehmann/Afum_Chr7_aneuploids.

## Acknowledgments

We thank the Duke University Sequencing and Genomic Technologies Core Facility for performing the sequencing and Dr. Paul Magwene, Dr. Adrianna San Roman, Dr. Marco Coelho, Dr. Vikas Yadav, and Anna Mackey for assistance and advice with analyzing the DNA- and RNA-sequencing data. We also thank Dr. Robert Cramer for providing the *A. fumigatus* strains and selectable marker plasmids and for offering invaluable insights about strain heterogeneity and antifungal drug resistance, Dr. Joe Vasselli for sharing a custom 3D-printed microscopy growth chamber, and the members of the Heitman Lab and the Duke mycology community for constructive discussion throughout this project. This study was supported by the NIH/NIAID grants 5R01AI172451-03 and 5R01AI170543-04 (JH) and NIH/NIGMS 1R35GM156119 (NPK). JH is also co-director and fellow of the Canadian Institute for Advanced Research (CIFAR) program Fungal Kingdom: Threats & Opportunities. Mass spectrometry data were obtained on equipment acquired with support of UW-Madison and grant 2024-70410-43730 (USDA).

## Author Contributions

Conceptualization: A.E.L. and J.H.; Investigation: A.E.L. and E.A.R.; Data analysis: A.E.L. and E.A.R.; Writing – original draft: A.E.L. and E.A.R.; Writing – review & editing: J.H. and N.P.K.; Funding acquisition: J.H. and N.P.K.

## Supplemental Information

Document S1: Figures S1-S9; Tables S2 and S4.

Table S1. High- and medium-impact coding region variants in all FK506-adapted isolates.

Table S3. Gene expression changes in Chr7 aneuploids +/- FK506.

Table S5. Sequence variants between our A1163 laboratory strain and the published reference genome.

## Materials and Methods

### Strains and growth conditions

The A1163 reference strain served as the haploid wild type in all genetic and transcriptomic comparisons presented in this study. Additional clinical and environmental isolates (Table S2), previously reported by Ross et al^86^ and Lofrgren et al^57^, were provided by Dr. Robert Cramer (Geisel School of Medicine at Dartmouth) and were also screened for aneuploidy in response to FK506. Strains were cultured on 1.5% agar Aspergillus Minimal Medium (AMM), which was prepared as described by Hill and Kafer^97^. All FK506 experiments on agar plates and in liquid culture were performed in AMM with FK506 at a concentration of 1 µg/mL or 0.5 µg/mL. Conidia were grown for experiments on AMM or AMM+FK506 for 3 days at 37°C. Unstable aneuploids were maintained on +FK506 plates to ensure retention of the ploidy differences. Conidia were collected from plates by flooding the surface of the plate with 0.01% Tween 80, manually dislodging conidia with a cell scraper, and filtering through Miracloth (Millipore Sigma). Conidia were counted with a hemacytometer.

### DNA extraction and sequencing

Biomass for DNA extraction was obtained by inoculating 5 mL AMM or AMM+FK506 with conidia at a density of 5x10^5^ conidia/mL and incubating at 37°C rotating at 70 rpm on a roller drum for 16-24 hours. Biomass was collected and washed 1x with diH_2_O, then frozen at -80°C for >1 hour and lyophilized until dry. DNA was extracted following a modified MasterPure (Biosearch Technologies) procedure. First, lyophilized tissue was bead-beaten with 2 mm glass beads for 1 minute. The remainder of the procedure was carried out according to the manufacturer’s instructions. Nucleic acid pellets were resuspended in nuclease-free water and quality was assessed by gel electrophoresis. DNA concentration was quantified with a Qubit fluorometer using the DNA broad-range kit (Thermo Fisher Scientific).

DNA sequencing was carried out by the Duke University Sequencing and Genomic Technologies Core Facility. Libraries were prepared using the Kapa HyperPrep kit (Kapa Bioscience). Pooled libraries were sequenced on an Illumina Novaseq X Plus instrument in 150 bp paired-end reads.

### DNA-seq data analysis

For sequence variant identification, DNA-seq reads were aligned to the A1163 reference genome^60^ with snippy^98^, using the default parameters. Resulting vcf files served as input for snpEff^99^, which compares variants to a preconfigured A1163 database and predicts variant effects on the coding transcript (low, medium, or high). For the FK506-adapted isolates, variants in coding regions only were identified with the flags –no-downstream, –no-upstream, and –no-intergenic. Unique variants were determined by filtering out any that were shared with our laboratory A1163 parental strain (Table S5).

Ploidy was determined by comparing coverage depth along 500 bp windows to genome-wide normalized read depth. Reads were aligned to the chromosome-level CEA10 reference assembly^100^ with minimap2^101^, using settings for paired-end short reads. Aligned reads were filtered by a threshold of q30 with samtools^102^. Mosdepth^103^ was used to calculate read depth over 500 bp windows. The resulting bed files were analyzed in R to compare depth in each window to genome-wide normalized coverage. Data were visualized with ggplot2^104^.

### Phenotypic assays

#### FK506 spot growth and imaging

10^5^ conidia were spotted on AMM + 1 µg/mL FK506 and incubated at 37°C for 2 days. Images of hyphal growth at the colony edge were acquired with an AxioScop 2 fluorescence microscope with 4x objective (Zeiss). Experiments were performed in at least 3 biologically independent replicates.

#### Imaging of submerged hyphal growth

100µL conidia at a concentration of 10^3^ conidia/mL in AMM ± 1 µg/mL FK506 were inoculated in a custom 3D-printed growth chamber^105^ encased between 2 transparent slides. Growth chambers were incubated at 37°C for 24 hours. Hyphal growth images were acquired on an AxioScop 2 fluorescence microscope with 10x and 20x objectives (Zeiss). Experiments were performed in biological triplicate.

### Transcriptomics

#### RNA extraction and sequencing

Biomass for RNA extraction was obtained from 10^6^ conidia/mL cultures in 25 mL AMM or AMM+FK506 grown for 16 hours at 37° with shaking at 200 RPM. Biomass was harvested, washed 1x with sterile diH_2_O, frozen at -80°C for >1 hour, and lyophilized until dry. Lyophilized tissue was bead-beaten for 1 minute with 2 mm glass beads and RNA was extracted with the Qiagen RNeasy Plant Mini Kit according to the manufacturer’s instructions (Qiagen). RNA concentration was measured a Qubit fluorometer using the RNA broad-range kit (Thermo Fisher Scientific).

RNA was sequenced by the Duke University Sequencing and Genomic Technologies Core Facility. Poly-A enriched libraries were prepared from samples with RIN ≥ 7 using the Kapa mRNA-seq HyperPrep kit (Kapa Bioscience). Pooled libraries were sequenced on an Illumina Novaseq X Plus instrument in 150 bp paired-end reads.

#### RNA-seq data analysis

Illumina universal adaptors were trimmed from paired reads with CutAdapt^106^ (v4.9). Trimmed reads were then aligned to the A1163 reference assembly^60^ with HISAT2^107^ (v2.2.1). Counts per gene were quantified with featureCounts^108^ (v2.0.6). Exploratory data analysis and differential expression analysis were performed with DESeq2^109^ (v1.44.0). Prior to analysis, transcript counts were normalized by transcripts per million (TPM) within each replicate and genes with at least 1 TPM in 1 sample were retained. Clustering and principal component analyses were performed using vst normalized values for the top 1000 most variable genes generated by DESeq2. Differential expression analysis was carried out in DESeq2 with alpha = 0.05 and a log_2_FC cutoff of +/- 1 was applied to identify DEGs from the resulting expression data. The R packages EnhancedVolcano (v1.22.0) (https://github.com/kevinblighe/EnhancedVolcano) and pheatmap (v1.0.13) (https://github.com/raivokolde/pheatmap) were used for data visualization.

#### nscR overexpression strain construction

The *nscR^OE^* strain was generated in the A1163 background by ectopic integration of *nscR* (AFUB_086680) driven by an *A. fumigatus gpdA* promoter amplified from the Af293 reference strain (Fungal Genetics Stock Center A1100). The *gpdA* promoter (_p_*gpdA*) was selected as the 996 bp immediately upstream of the *gpdA* (Afu5g01970) open reading frame using primers AEL68 and AEL69. The promoter was fused to the *nscR* open reading frame and the 504 bases immediately 3’ of the stop codon, which was amplified from A1163 gDNA with primers AEL70 and AEL71, and the hygromycin resistance marker *hygR*. The _p_*gpdA^AF^-nscR* overexpression construct fused to *hygR* selectable marker was amplified with primers AEL75 and AEL76. The resulting product was used to transform A1163 protoplasts as described previously^110,111^. Transformants were selected on osmotically-stabilized minimal medium (AMM + 1.2 M sorbitol) containing hygromycin. Successful transformants were single spored, and genomic integration of the construct was verified by PCR and expression was measured by RT-qPCR as described below, with RNA obtained from WT and *nscR^OE^* under the same conditions ± FK506 used for RNA-seq. All primer sequences used are listed in Table S5.

#### Reverse transcriptase quantitative PCR (RT-qPCR)

2.5 to 5 µg of RNA was DNase treated using the TURBO DNA-free kit (Invitrogen) following the manufacturer’s instructions. Post-DNase treatment RNA integrity was verified by running a sample of the RNA on an agarose gel. cDNA synthesis was carried out using 500 ng of DNase-treated RNA as described previously^104^. RT-qPCR was run on a QuantStudio 3 or QuantStudio 6 Pro system (Thermo Fisher Scientific). Gene expression was normalized to *actA* and *tubA* in all experiments. Fold change values were calculated by the 2^-ΔΔCt^ method^112^.

#### Metabolomic studies

Four biological replicates of each strain were cultured at 10^6^ conidia/mL in 25 mL of GMM medium supplemented with 1 µg FK506/mL and incubated for two days at 37 °C with shaking at 200 rpm. After the fermentation period, each fungal culture and four medium replicates (considered as blanks) were extracted by maceration with 25 mL of EtOAc/MeOH/AcOH (89:10:1) overnight, with shaking at 120 rpm. The organic phase was then separated and evaporated to dryness under reduced pressure. The resulting organic extracts were dissolved in MeOH at a concentration of 50 mg/mL, and serial dilutions (1:10 and 1:50) were prepared to obtain solutions at 0.1 mg/mL for LC-MS analysis. Then, a 5 µL aliquot of each sample was analyzed by UPLC coupled to an Orbitrap Exploris™ 240 mass spectrometer, acquiring both MS^1^ and MS/MS spectra, using an Acquity BEH-C18 column (2.1 mm × 100 mm, 1.7 μm).

The chromatographic method consisted of a one minute isocratic elution of 10% MeCN in H_2_O containing 0.05% formic acid, followed by a linear gradient from 10% to 100% MeCN over 20 minutes, and a 2.5-minute wash with 100% MeCN. The flow rate was maintained at 0.3 mL/min throughout the run, with alternating positive and negative ion detection over an *m*/*z* range of 120–1500 for MS/MS acquisition. Precursor ions were fragmented using higher-energy collisional dissociation (HCD) with dynamic exclusion set to 3 s.

Once the LC-MS/MS data were acquired, the raw files generated from these analyses were converted to .mzML format using MSConvert^113^, applying the peak-picking filter for both MS^1^ and MS^2^. Spectra were processed in MZmine^114^ (v4.7.8). Mass detection was performed using the lowest signal factor, with a noise threshold set at 5.0 for MS^1^ and 2.5 for MS^2^. A chromatogram was constructed using the Chromatogram Builder algorithm in positive polarity mode, with parameters set to a minimum of 4 consecutive scans, a minimum intensity of 1×10^4^ for consecutive scans, a minimum absolute height of 5×10^4^, and a mass tolerance of 10 ppm.

The resulting chromatogram features were then processed using Savitzky-Golay smoothing (5 scans retention time width), the Local Minimum Feature Resolver, a 13C isotope filter, the Join Aligner, Feature Finder, and duplicate peak filter. From this processed feature list, features detected in the blank replicates were manually removed, yielding a total of 4,495 features. This curated feature list was used for principal component analysis (PCA) with Pareto scaling and missing value imputation (set at 1/5 of the minimum value), for pairwise comparisons using unpaired Volcano Plots, and for molecular network construction following the Feature-based Molecular Networking workflow with default parameters^115^.

The abundance of secondary metabolites was represented using the area under the curve of the target feature (included in chromatograms, bar graphs, and heatmaps), based on LC-MS data acquired in either negative or positive ionization mode. PCA plots, Volcano Plots, chromatograms, bar graphs, and heatmaps were generated with Prism 10.6.0. For the heatmap, z-scores were calculated based on the average abundance of each metabolite across the different strains. Metabolites that were not detected in any strain were labeled as undetected. The molecular network was visualized using Cytoscape^116^ (v3.10.3). The identities of fumitremorgin C, fumagillin, pyripyropene A, fumiquinazoline F, neosartoricin, and FK506 were confirmed by comparison of their chromatographic and spectrometric data with authentic standards. The identity of pseurotin A was supported by matching the fragmentation spectrum of the detected molecular ion to a high-quality gold-standard reference deposited in the GNPS database (https://gnps.ucsd.edu/ProteoSAFe/gnpslibraryspectrum.jsp?SpectrumID=CCMSLIB00011906568#%7B%7D). The ergosterol-type triterpenoids class was supported by the predicted formula and by the comparison of the fragmentation pattern with those reported by Lucie Ory et al^117^.

#### Antifungal susceptibility tests

Broth dilution MIC assays were performed according to the established standard CLSI procedure^118^. MIC tests were performed in 3 to 4 biological replicates.

#### Quantification of cyp51A and cyp51B transcript abundances

RNA was obtained from two distinct culture conditions to provide for potential condition-specific, rather than drug-specific, effects on transcript abundances. In the first condition, 2.5x10^5^ conidia were inoculated in 5 mL RPMI medium and incubated at 37°C rotating at 70 rpm. At 22 hours, medium was removed and replaced with 5 mL RPMI + 0.2 µg/mL voriconazole (0.1 µL of a 10 mg/mL stock in DMSO) or 0.1 µL DMSO. Cultures were incubated under the same conditions for an additional 2 hours. For the second condition, 10^6^ conidia/mL were inoculated in AMM with 0.1 µg/mL voriconazole or DMSO and incubated at 37°C with shaking at 200 rpm. Biomass was collected at 24 hours. For both conditions, RNA was extracted and cDNA synthesized as described above. qPCR was carried out as described for the *nscR^OE^*strain, and *cyp51A* and *cyp51B* transcript abundances was calculated relative to *actA*. All qPCR primers used in this study are listed in Table S4.

