## Supplementary material for "Aneuploidy promotes transient stress adaptation and metabolic flexibility in the human fungal pathogen *Aspergillus fumigatus*": Document S1

| Strain | Clade | Source | Geographic origin | *MAT* |
|---|---|---|---|---|
| AF100-1_18 | 1 | Clinical | USA | *1-1/1-2* |
| A_fum_Moose_3_1_2 | 1 | Environmental | USA | *1-1* |
| AF100-12_5 | 1 | Clinical | USA | *1-1/1-2* |
| AF100-1_3 | 1 | Clinical | USA | *1-1/1-2* |
| A_fum_Oryx_1 | 1 | Environmental | USA | *1-2* |
| AF100-12_8 | 1 | Clinical | USA | *1-2* |
| 08-19-02-10 | 2 | Environmental | Netherlands | *1-2* |
| 10-01-02-27 | 2 | Clinical | United Kingdom | *1-2* |
| F15390 | 2 | Clinical | United Kingdom | *1-1* |
| W72310 | 3 | Clinical | USA | *1-2* |

**Table S2. Characteristics of 10 genetically-diverse *A. fumigatus* strains.** Strain metadata adapted from: Lofgren LA, Ross BS, Cramer RA, Stajich JE (2022) The pan-genome of *Aspergillus fumigatus* provides a high-resolution view of its population structure revealing high levels of lineage-specific diversity driven by recombination. PLOS Biol 20(11): e3001890. https://doi.org/10.1371/journal.pbio.3001890.

| ID | Sequence | Description |
|---|---|---|
| AEL68 | AGATATACGGCGTATGATTCAGAGC | Amplify *gpdA* promoter from AF293 gDNA; ligate promoter to AFUB_086680. |
| AEL69 | TGTGTAGATTCGTCTGGTACTGAGC | Amplify *gpdA* promoter from AF293 gDNA. |
| AEL74 | GTTTCCGCCGATTACTTTTCTCCATTGTGTAGATTCGTCTGGTACTGAGC | Add microhomology to AFUB_086680 5' for plasmid construction (with AEL68). |
| AEL70 | ATGGAGAAAAGTAATCGGCGG | Amplify AFUB_086680 from A1163 gDNA. |
| AEL71 | GGTTTTGTTGTGTTTTAGTCCAAGG | Amplify AFUB_086680 from A1163 gDNA; ligate promoter to AFUB_086680. |
| AEL72 | CCTTGGACTAAAACACAACAAAACCATGGGGAAGGCCATCCAGCC | Amplify hygR cassette and split origin from plasmid BS311. |
| AEL80 | GTCGTGTCTTACCGGGTTGG | Amplify hygR cassette and split origin from plasmid BS311. |
| AEL81 | TGAGTCCAACCCGGTAAGAC | Split origin and amplify other half of BS311 backbone. |
| AEL73 | GCTCTGAATCATACGCCGTATATCTTCTCGCTTCCGGCGGCATCG | Split origin and amplify other half of BS311 backbone. |
| AEL75 | GACCCTTCCTTTTGTAGGTTATTACCCTGCCCTTGGGTATCGGCGTATTGGGTGTTACGGAGC | Amplify *p.gpdA-nscR-hygR* construct from pAEL5. |
| AEL76 | GGGTAGGTCTCCTTCCGGAGGCCAACGATTTTGTTGTGGTAGATATACGGCGTATGATTCAGAGC | Amplify *p.gpdA-nscR-hygR* construct from pAEL5. |
| AEL86 | TGAATGTCGGGCTCGACCTTGC | *nscR* qPCR fwd. |
| AEL87 | CCAACCATTGAGGCGTGCTACC | *nscR* qPCR rvs. |
| AEL88 | CCGAACAGAACCGCCAATGG | *cyp51A* qPCR fwd. |
| AEL89 | AACGCCCAGGTAGACTGTGG | *cyp51A* qPCR rvs. |
| AEL90 | GCAAGCTGCGAGATGTTTGC | *cyp51B* qPCR fwd. |
| AEL91 | AGCGCATCAGAGGTGAGACC | *cyp51B* qPCR rvs. |
| 53313 | GTGATGTCGACGTCCGTAAGG | *actA* qPCR fwd. |
| 53314 | GCATACGGTCGGAGATACCG | *actA* qPCR rvs. |

**Table S4. Oligonucleotides used in this study.**

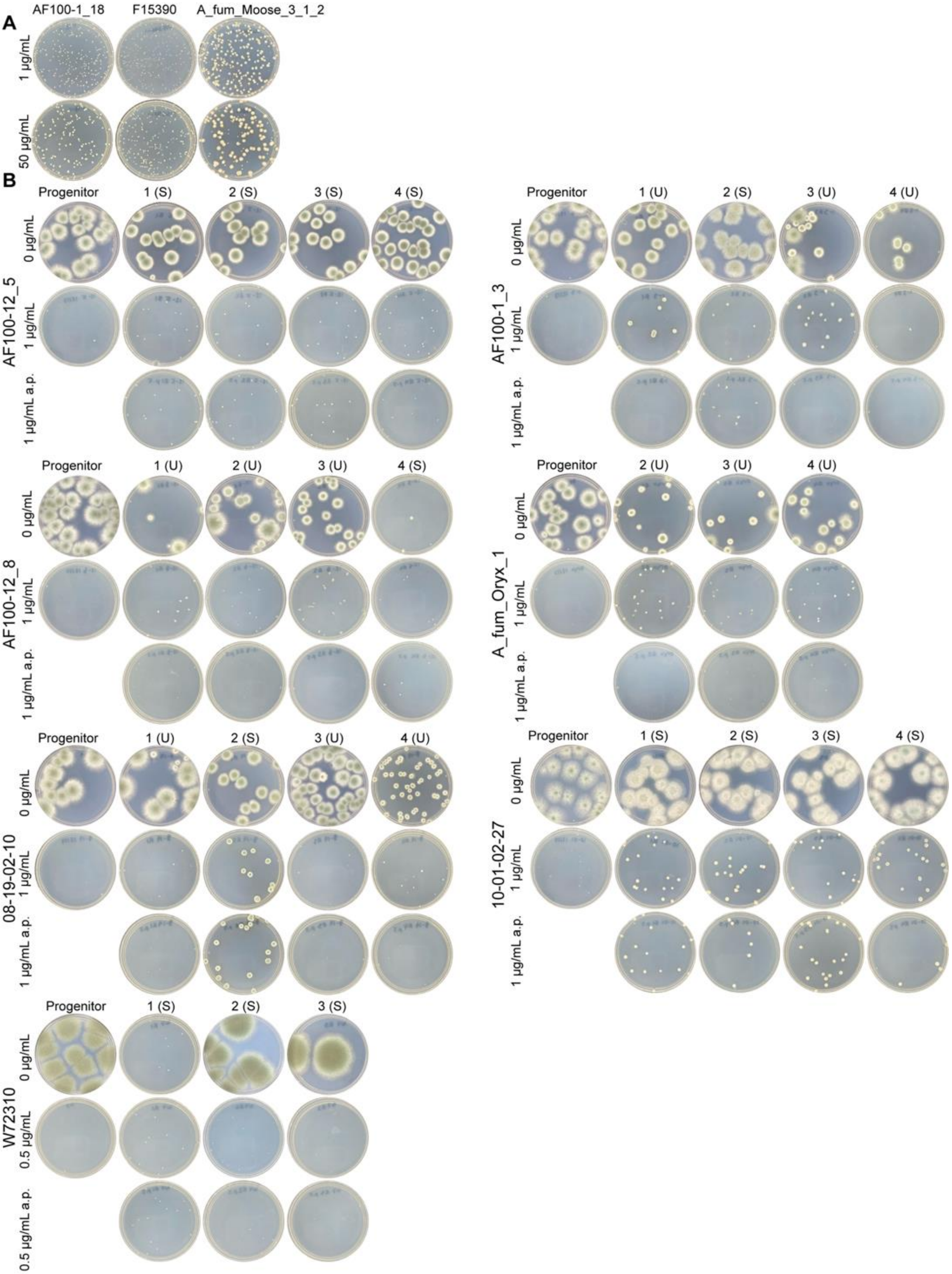
A
AF100-1_18
F15390
A_fum_Moose_3_1_2
1 μg/mL
50 μg/mL
B
AF100-12_5
Progenitor
1 (S)
2 (S)
3 (S)
4 (S)
0 μg/mL
1 μg/mL
1 μg/mL a.p.
AF100-1_3
Progenitor
1 (U)
2 (S)
3 (U)
4 (U)
0 μg/mL
1 μg/mL
1 μg/mL a.p.
AF100-12_8
Progenitor
1 (U)
2 (U)
3 (U)
4 (S)
0 μg/mL
1 μg/mL
1 μg/mL a.p.
A_fum_Oryx_1
Progenitor
2 (U)
3 (U)
4 (U)
0 μg/mL
1 μg/mL
1 μg/mL a.p.
08-19-02-10
Progenitor
1 (U)
2 (S)
3 (U)
4 (U)
0 μg/mL
1 μg/mL
1 μg/mL a.p.
10-01-02-27
Progenitor
1 (S)
2 (S)
3 (S)
4 (S)
0 μg/mL
1 μg/mL
1 μg/mL a.p.
W72310
Progenitor
1 (S)
2 (S)
3 (S)
0 μg/mL
0.5 μg/mL
0.5 μg/mL a.p.

**Figure S1. Phenotypes of FK506-adapted isolates in diverse strain backgrounds.** (*A*) 3/10 isolates exhibit robust growth on 1 µg/mL FK506 (top) and 50 µg/mL FK506 (bottom). (*B*) Phenotypes of FK506-adapted isolates in 7 strain backgrounds before and after passage on drug-free medium (a.p.). Isolates are characterized as stable (S) if they exhibited the same enhanced growth phenotype on FK506 after 15 days of drug-free passage, and unstable (U) if the phenotype was lost after passage. With the exception of W72310, images were taken after 3 days of growth at 37°C. W72310 has a slow rate of growth and images were taken after 5 days of growth at 37°C.

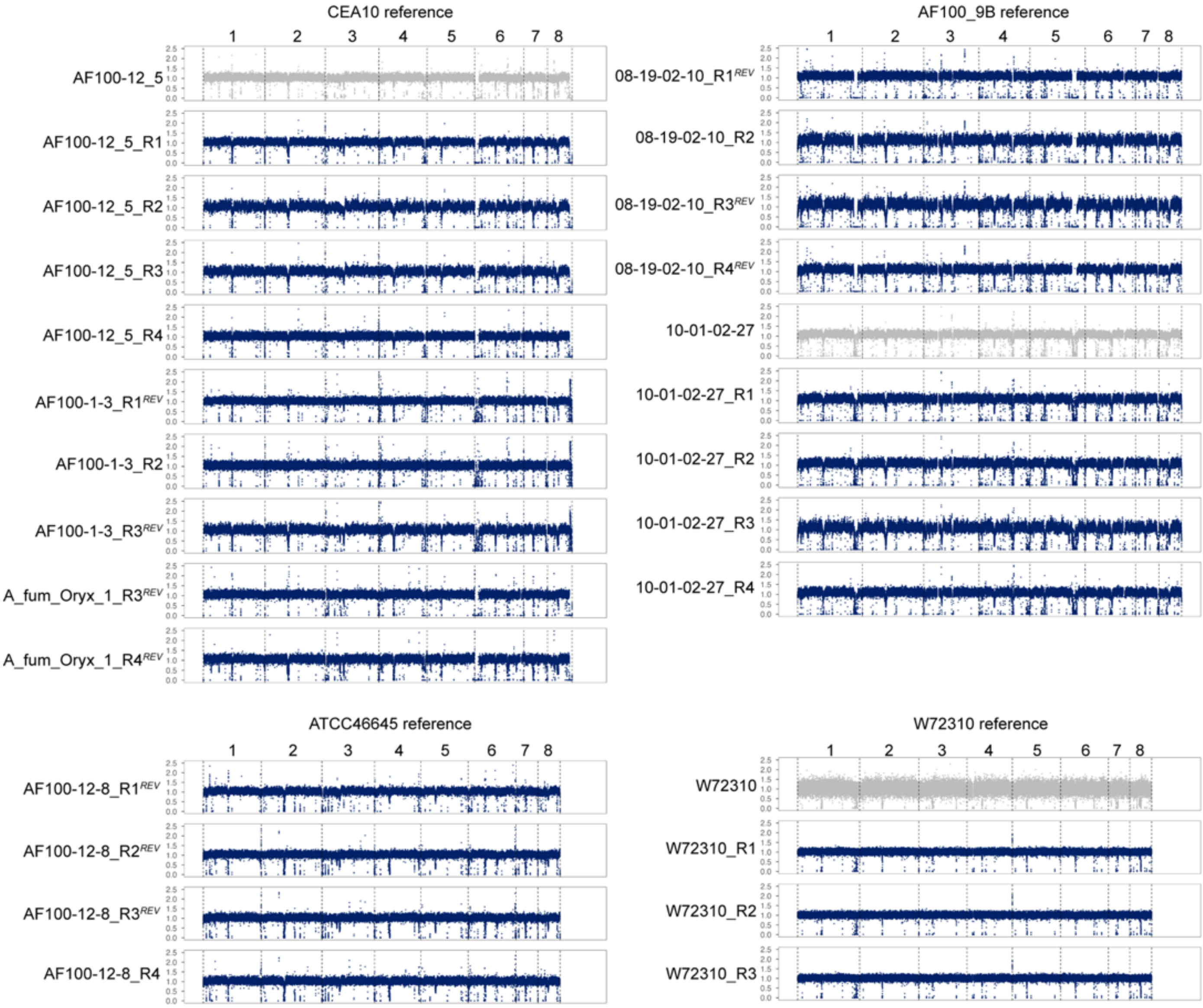


**Figure S2. Coverage plots of parental, reverted, and stable FK506-adapted isolates.** Additional coverage plots for isolates not shown in Fig. 2B. Reads were mapped to a closely-related assembly as indicated above each set of plots.

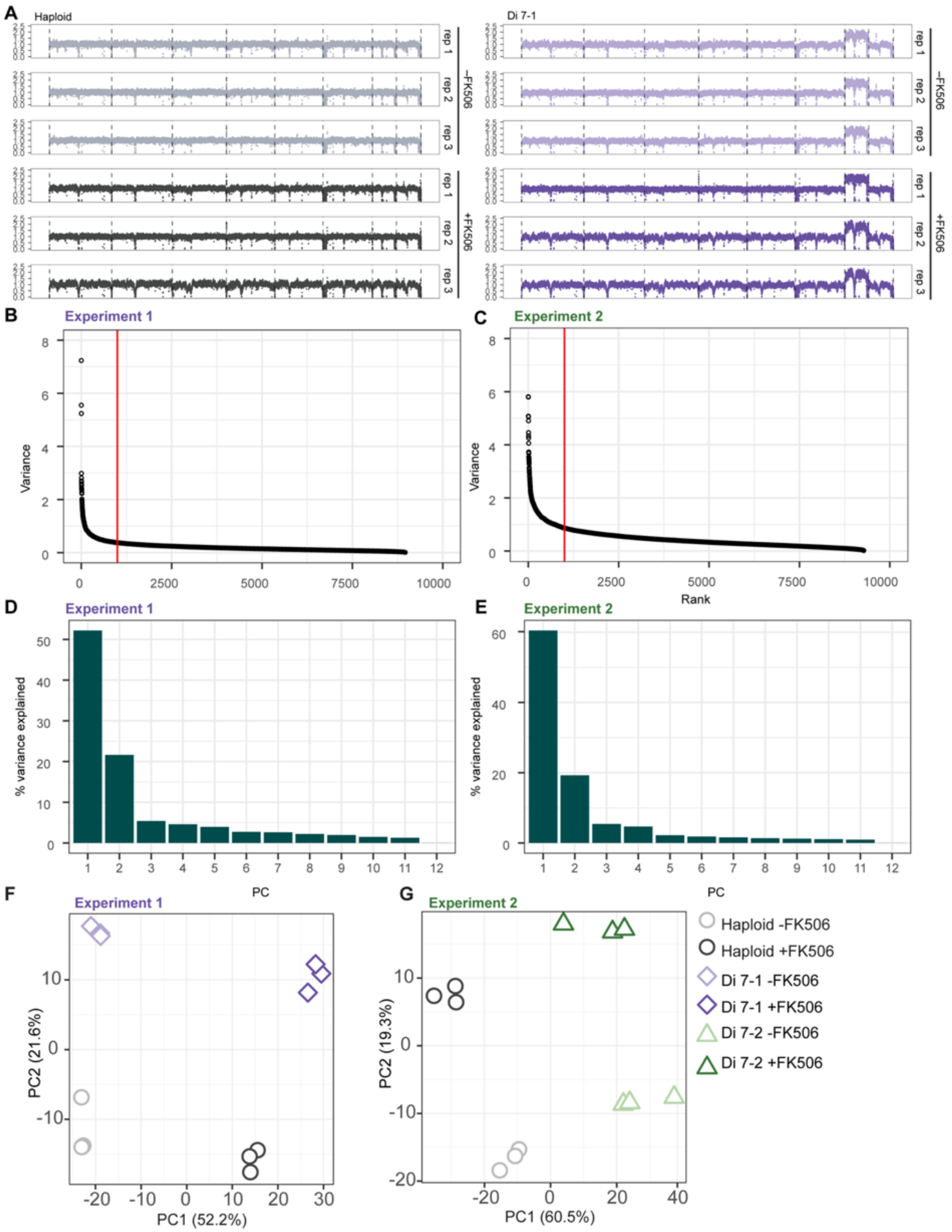


**Figure S3. RNA-seq exploratory data analysis.** (*A*) WGS coverage plots from DNA collected in parallel with RNA in Experiment 1 (Di 7-1). Genome-wide normalized read depth, calculated over 500 bp intervals, is plotted relative to the CEA10 reference assembly coordinates. (*B* and *C*) TPM-filtered genes were ordered by vst-normalized expression between conditions to identify the top 1000 most variable genes to use for principal component analysis in experiment 1 (*B*; Di 7-1) and 2 (*C*; Di 7-2). (*D* and *E*) % variance captured by each principal component in PCA of the top 1000 most variable genes in experiment 1 (*D*) and 2 (*E*). (*F* and *G*) PCA of the top 1000 most variable genes in Experiment 1 (*F*) and Experiment 2 (*G*).

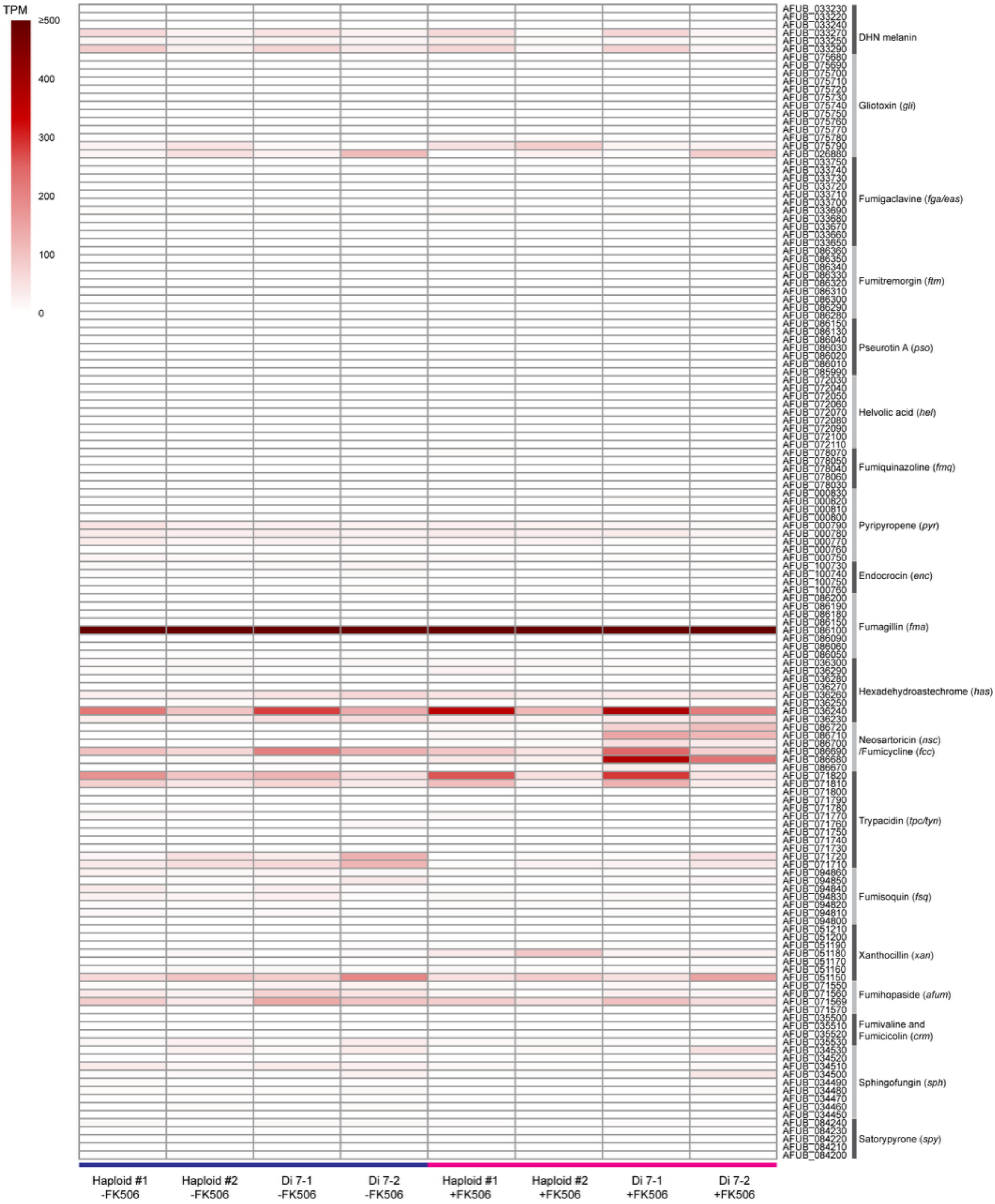


**Figure S4. Secondary metabolite BGC transcription in A1163 haploids and Chr7 disomic strains (Di 7) +/- FK506.** Average TPM from 3 technical replicates plotted for A1163 BGCs.

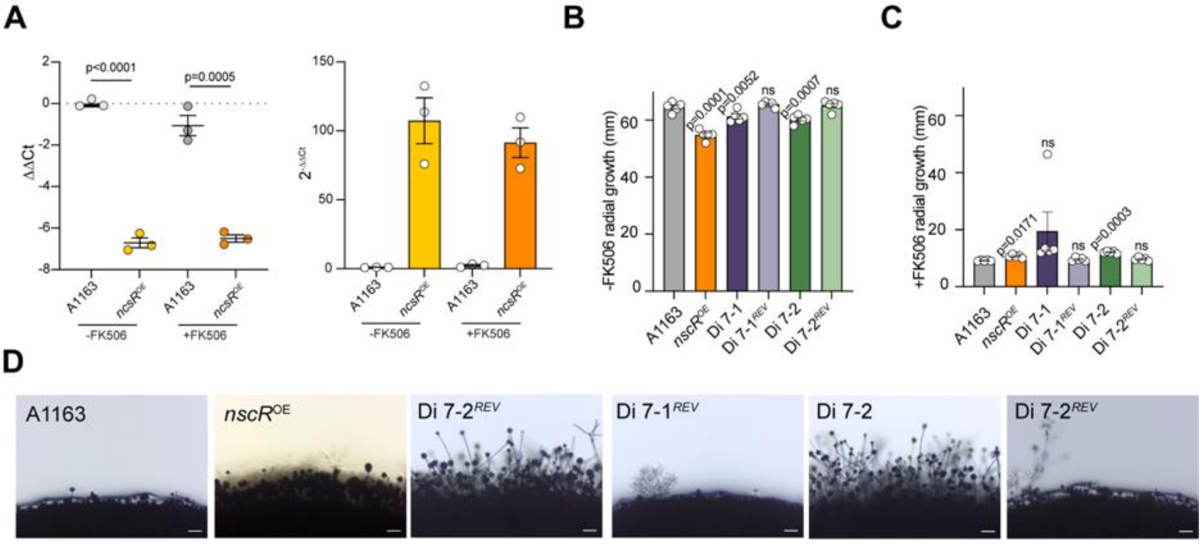


**Figure S5. Additional *nscR$^{OE}$* and Chr7 aneuploid phenotypes.** (*A*) Fold change values of *nscR* transcript in WT and *nscR*$^{OE}$ strains +/-FK506 relative to WT untreated. RNA was extracted from 16-hour cultures seeded at $10^6$ conidia/mL in liquid AMM +/- 1 µg/mL FK506 and grown at 37°C with shaking at 200 rpm. ΔCt values were calculated by normalizing to *actA* and ΔΔCt values relative to WT untreated. Points indicate biological replicates and are represented as averages of 3 technical replicates; error bars indicate SEM. Statistical test between wild-type and *nscR$^{OE}$* in each condition is an unpaired t-test. (*B* and *C*) $10^4$ conidia were point-inoculated on AMM ±FK506 and grown at 37°C for 4 days. Radial growth measurements representative of n = 5 biologically independent replicates; statistical tests are one-way ANOVA with Dunnett's multiple comparison test. (*D*) $10^5$ conidia were spotted on AMM + 1 µg/mL FK506 and grown at 37°C for 2 days. Microscopy images of hyphal and conidiophore growth at the colony periphery are representative of n = 3 biologically independent replicates. Scale bars indicate 100 µm.

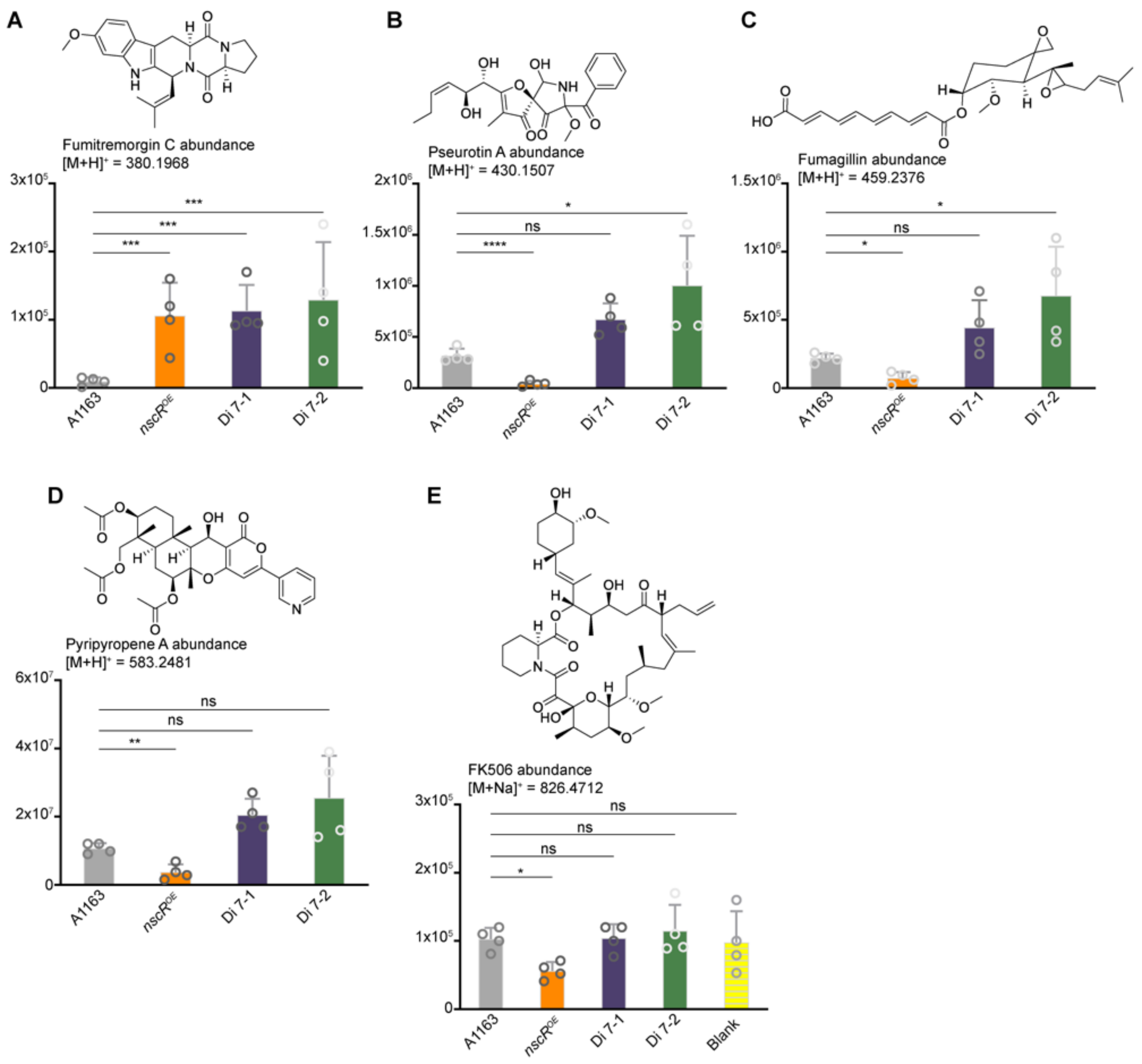


**Figure S6. Selected secondary metabolites overproduced in the aneuploids compared to the haploid strain, and FK506 abundance**. Abundance of fumitremorgin C (*A*), pseurotin A (*B*), fumagillin (*C*), pyripyropene A (*D*), and FK506 (*E*) in fungal cultures of haploid, *nscR*$^{OE}$, Di 7-1, and Di 7-2, represented as bar graphs showing the average area under the curve of the respective molecular ion intensities. One-way ANOVA followed by Dunnett's post-hoc test was applied on log-transformed data to compare *nscR*$^{OE}$ and Di-7 strains to A1163 haploid. * = $p < 0.05$; ** = $p < 0.01$; *** = $p < 0.001$; **** = $p < 0.0001$.

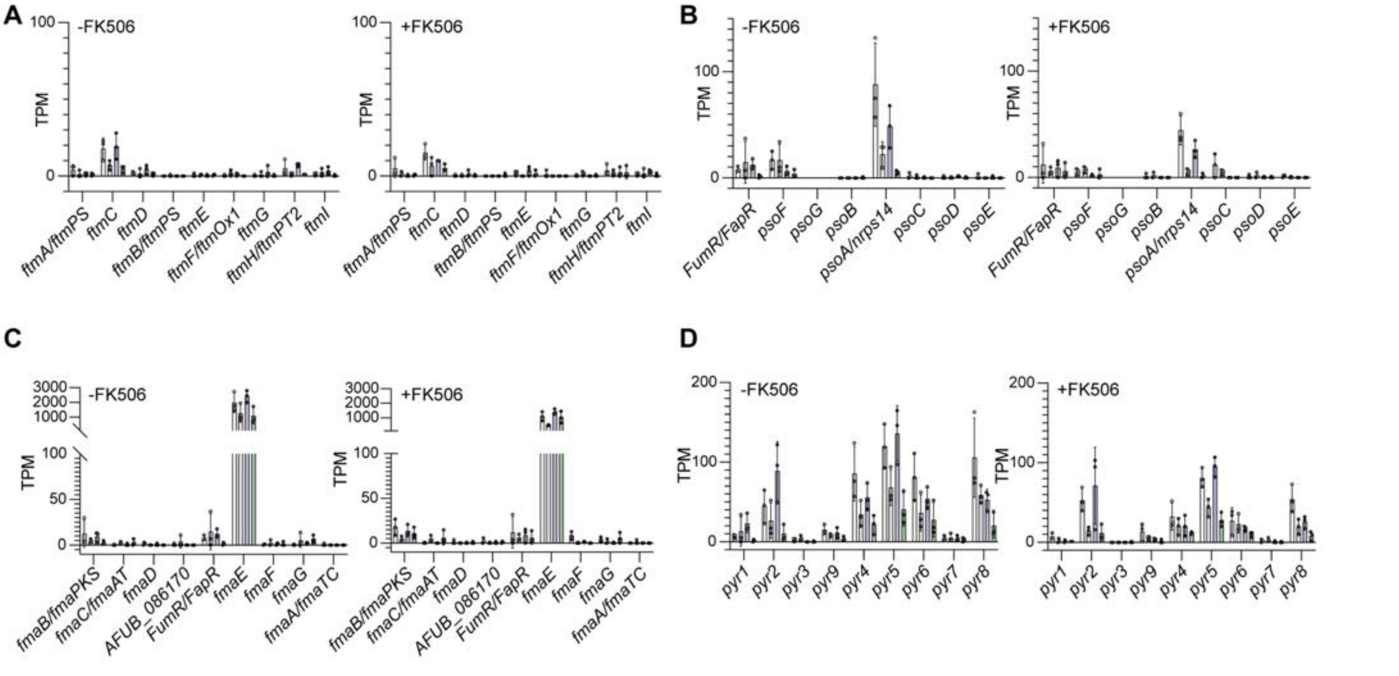


**Figure S7. Transcript abundances of genes in selected secondary metabolite BGCs.** Transcript abundances are represented as transcripts per million (TPM) for fumitremorgin C (*A*), pseurotin (*B*), fumagillin (*C*), and pyripyropene A (*D*) BGCs. Points represent technical replicates; error bars indicate standard deviation. In each plot 2 independent RNA-seq experiments are represented; light grey and purple bars indicate haploid and Di 7-1, respectively, from experiment 1 and dark grey and green bars are haploid and Di 7-2 from experiment 2.

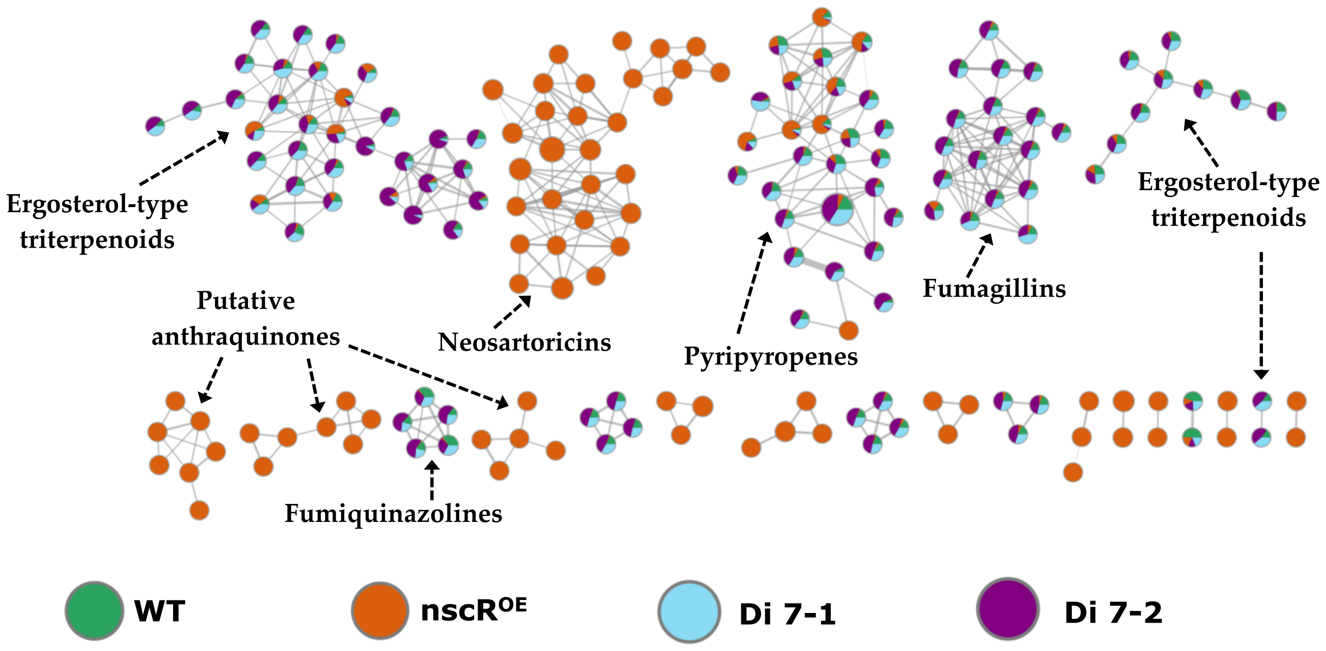


**Figure S8. Segment of the molecular network generated by FBMN (in positive mode), showing the main classes of secondary metabolites that define the characteristic metabolomic profile of WT, *nscR$^{OE}$*, Di 7-1, and Di 7-2 strains.** Distinct ergosterol-type triterpenoids, pyripyropenes, fumagillins, and fumiquinazolines derivatives are predominantly present in the aneuploid strains, whereas neosartoricin derivatives are exclusively detected in the *nscR$^{OE}$* strain.

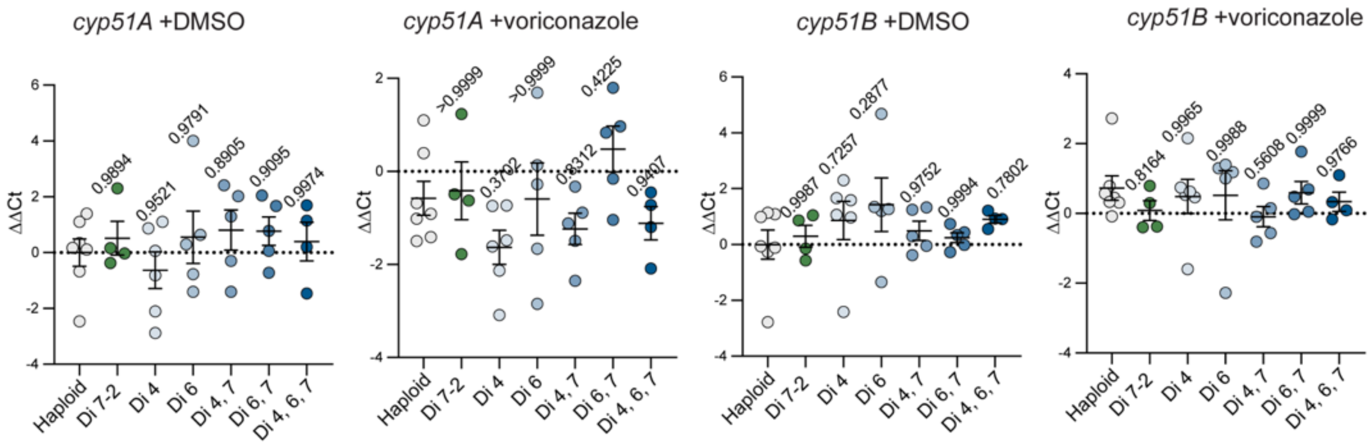


**Figure S9. *cyp51A/B* qPCR results from AMM shaking flask cultures.** $10^6$ conidia/mL were inoculated in 25 mL AMM +DMSO or +0.1 µg/mL voriconazole and incubated at 37° with shaking at 200 rpm. Biomass was collected for RNA extraction at 24 hours. qPCR was performed to assess the relative abundances of *cyp51A* and *cyp51B* transcripts. ∆Ct values were calculated relative to *actA* and ∆∆Ct values are relative to haploid +DMSO. Statistical significance was assessed by one-way ANOVA with Dunnett's multiple comparison test. Points represent averages of technical triplicates from n = 4-7 biological replicates.
